# The effect of oxidative stress on the Adenosine A2a Receptor structure, activity and signalling

**DOI:** 10.1101/2024.10.31.620957

**Authors:** Idoia Company-Marin, Joseph Gunner, David Poyner, John Simms, Andrew R. Pitt, Corinne M. Spickett

## Abstract

The adenosine 2a receptor (A2aR) is a G-protein coupled receptor that has important anti-inflammatory effects in response to some agonists and consequently is considered a therapeutic target. Its activity is affected by local membrane lipid environment and presence of certain phospholipid classes, so studies should be conducted using extraction methods such as styrene maleic acid co-polymers (SMA) that retain the local lipids. Currently, little is known about the effect of oxidative stress, which may arise from inflammation, on the A2aR. Therefore it was over-expressed in *Pichia pastoris*, SMA was used to extract the A2aR from cell membranes and its response to ligands was tested in the presence or absence of the radical initiator AAPH or reactive aldehyde acrolein. SMA-extracted A2aR was able to undergo conformational changes, measured by tryptophan fluorescence, in response to its ligands but oxidative treatments had no effect on the structural changes. Similarly, the treatments did not affect temperature-dependent protein unfolding. In contrast, in HEK293 cells expressing the A2aR, oxidative treatments increased cAMP levels in response to the agonist NECA, independently of adenylate cyclase activity. Thus, oxidative stress may be a homeostatic mechanism that abrogates inflammation via the A2aR signalling pathway. (max. 200 words - 194)

## 1. Introduction

Adenosine receptors belong to the largest class of G-protein coupled receptors (GPCRs), the rhodopsin-like receptor family, and are characterised as being activated by extracellular adenosine as a natural agonist, and inhibited by purine base xanthines, such as caffeine and theophylline (Goulding et al., 2018). The adenosine A2a receptor (A2aR) is an important GPCR that binds adenosine at nanomolar concentrations to inhibit inflammation and platelet aggregation. Other effects include vasodilation of coronary arteries to increase myocardial blood flow, causing hypotension, and inhibiting dopaminergic activity in the brain (Bruzzese et al., 2020). The downstream actions of the A2aR depend on its interaction with cytosolic G proteins, typically the heterotrimeric Gs protein, leading to activation of adenylate cyclase and production of cAMP to activate protein kinase A (PKA). This initiates a cascade of phosphorylation, although the specific signalling pathways are tissue and cell type dependent (de Lera Ruiz et al., 2014). There has been considerable interest in the A2aR as a therapeutic target, as several studies have demonstrated the anti-inflammatory and immunosuppressive roles of various A2aR agonists (Castro et al., 2020, El-Shamarka et al., 2020, Ikram et al., 2020, Leone and Emens, 2018, Li et al., 2018, Wang et al., 2022, Wang et al., 2019).

Inflammatory conditions lead to redox imbalance and oxidative stress through activation of innate immune cells and release of reactive oxygen species, which cause oxidative damage to a variety of biomolecules (Egea et al., 2018). Polyunsaturated phospholipids are susceptible to peroxidation and subsequent rearrangement or breakdown of the oxidized fatty acyl chain to yield electrophilic lipid peroxidation products such as phospholipids containing cyclopentenone rings or small reactive molecules including 4-hydroxynonenal, 4-hydroxyhexanal, malondialdehyde and acrolein (Sousa et al., 2017, Reis and Spickett, 2012, Vigor et al., 2014). As well as altering membrane phospholipid composition, the electrophilic lipid peroxidation products can react covalently with proteins to form adducts, a process known as lipoxidation (Aldini et al., 2015, Domingues et al., 2013, Spickett and Pitt, 2020). There is substantial evidence that lipoxidation can alter protein conformation, protein-protein interactions and enzymatic activity, with both inhibitory and activation effects reported, as reviewed by (Viedma-Poyatos et al., 2021). For example, acrolein, hydroxyhexanal and malondialdehyde inhibit the activity of the glycolytic enzyme pyruvate kinase (Sousa et al., 2019), acrolein and hydroxynonenal inhibit the dual specificity phosphatase PTEN (Covey et al., 2010), while H-ras (Oeste et al., 2011, Oliva et al., 2003) and the membrane channel TRPA1 (Takahashi et al., 2008) can be activated by electrophilic prostaglandins. Despite the important role of the A2aR in limiting inflammation, there have been few studies on the effects of oxidative damage and lipoxidation on its activity and downstream signalling, representing a significant gap in understanding.

The A2aR exists in dynamic equilibrium between the “inactive” conformations (absence of agonist or presence of an antagonist), intermediate-active (achieved on binding an agonist), and fully active states when it binds both an agonist and a cytosolic protein (e.g. Gs) to activate downstream signalling pathways (Manglik and Kruse, 2017, Weis and Kobilka, 2018).

Agonists, which include the synthetic compounds 5′-(N-Ethylcarboxamido) adenosine (NECA) (Lebon et al., 2011) and UK-432097 (Xu et al., 2011), bind in the orthosteric binding pocket of the A2aR, which is highly conserved and on the extracellular part of the receptor (Steyaert and Kobilka, 2011). This triggers multiple conformational changes, transduced from the orthosteric pocket to the intracellular ends of transmembrane helixes TM3, TM5, TM6, and TM7, which are essential for interaction with the cytosolic Gs protein and further stabilize the active conformation (Venkatakrishnan et al., 2013). Conformational changes in response to different agonists and antagonists can be studied *in vitro* using tryptophan fluorescence, as its fluorescence wavelength and intensity are strongly influenced by the polarity of its microenvironment and non-covalent interactions such as hydrogen bonding (Ghisaidoobe and Chung, 2014). Tryptophan (trp) emits at shorter wavelengths (λEM 330-340 nm) when buried in the hydrophobic core of a protein in its ground state, whereas it emits at longer wavelengths (340- 355 nm) when exposed to water in its excited state (Burstein et al., 1973). The native A2AR contains seven Trp residues, in TM1, TM4, TM6 and TM7 (Lebon et al., 2011), so its activation state is amenable to study by this approach (Routledge et al., 2020).

Many previous studies of the A2aR activation were conducted in vitro using detergent-extracted proteins. However, it is known that GPCRs, including the A2aR, can be affected by the local membrane lipid environment and the biophysical properties of the membrane, such as fluidity, thickness, curvature and lateral pressure (Baccouch et al., 2022). These properties are largely determined by the membrane lipid composition: anionic phospholipids phosphatidylserine (PS), phosphatidylinositol (PI), and phosphatidylglycerol (PG) have been suggested to act as positive regulators of GPCRs through close interactions, whereas phosphatidylethanolamine (PE) acts as a negative regulator, and phosphatidylcholine (PC) has a neutral effect (Dawaliby et al., 2016, Jones et al., 2020). Cholesterol affects the bilayer fluidity and can regulate ligand binding affinity via either direct or indirect allosteric mechanisms (Geiger et al., 2021, Paila and Chattopadhyay, 2009).

Consequently, studies of A2aR activity and mechanism should be conducted in the present of native membrane lipids and under conditions where bulk membrane properties such as lateral pressure are retained. This can be achieved by the use of membrane-protein extraction copolymers such as styrene maleic acid (SMA), which inserts into the membrane and extracts membrane proteins within small discs of bilayer styrene maleic acid encircled by the polymer, referred to as synthetic nanodiscs or SMALPs (SMA lipid particles) (Jamshad et al., 2015a, Knowles et al., 2009). As SMALPs retain the native lipid environment of the membrane protein they offer a more physiological model system, and many papers have reported improved native structure and activity by this method (Jamshad et al., 2015b, Logez et al., 2016). An alternative co-polymer is diisobutylene maleic acid (DIBMA), which has the advantage that it has been reported to cause less perturbation of phospholipid bilayer dynamics than SMA (Real Hernandez and Levental, 2023).

To address the question of the effect of oxidative stress on the structure and function of the A2aR, two complementary approaches were used. Firstly, a his-tagged construct of the human A2aR was over-expressed in *Pichia pastoris* and extracted from membranes with SMA2000 or alternative polymers. The nanodisc-encapsulated A2aR was subjected to oxidative treatments and conformational changes and protein stability were monitored by tryptophan fluorescence and thermal unfolding measurements, respectively. Secondly, the A2aR was transiently over- expressed in HEK293 cells, and the effects of oxidative stress on ligand binding and cellular signaling were monitored by measuring cAMP production. The combined information provides novel insight into the effect of oxidative stress conditions on the A2aR and downstream signalling.

## 1. Materials and Methods

### 1.1. Materials

The de-glycosylated C-terminal truncated A2aR (A316) with multiple tags including decahistidine, FLAG, and biotin in pPICZB developed by Jamshad and optimized by Bawa (Jamshad et al., 2015a) was utilized for expression in *Pichia pastoris*, essentially following their procedures. HisPur™ Ni-NTA Resin was obtained from Thermo Fisher Scientific, as was 7- Diethylamino-3-(4’-Maleimidylphenyl)-4-Methylcoumarin (CPM) (Invitrogen™ D346) and CPM stock was prepared at 5 mg/mL. ZM241385 and 2,2’-azobis(2-amidinopropane) dihydrochloride were purchased from Merck.

Dulbecco′s Phosphate Buffered Saline (DPBS) was obtained from Sigma-Aldrich, UK (D8537).

### 1.2. Expression of the A2AR in Pichia pastoris

*P. pastoris* SMD1163 cells were transformed with the pPICZB-hA2aR(A316) plasmid and grown in the presence of 100 µg/ml zeocin, as described previously (Jamshad et al., 2015a). After initial growth at 30°C in buffered glycerol complex medium (BMGY) (1% yeast extract, 2% peptone, 100 mM potassium phosphate at pH 6.0, 1,34% yeast nitrogen base without amino acids, 0.00004% biotin, 1% glycerol) also containing zeocin, cells were washed and transferred to buffered methanol complex medium (BMMY) where the glycerol was replaced with methanol and grown at 22°C for 24 h. After harvesting pellets were either used immediately for cell membrane preparation and analysis of protein expression, or frozen at -80°C for subsequent experiments.

### 1.3. P. pastoris membrane preparation

*P. pastoris* cells expressing the A2aR were resuspended in 50 mM sodium phosphate pH 7.4, 100 mM NaCl, 2 mM EDTA, 5% glycerol and 1 mM phenylmethylsulfonyl fluoride to a final concentration of 0.3 g/mL. Lysis was achieved by addition of an equal volume of acid-washed glass beads (425-600 µm) with 10 cycles of vortexing for 30 s followed by 1 min on ice. Unbroken cells and large debris were removed by centrifugation at 4000 g for 5 min at 4°C and 10,000 g for 10 min at 4°C. The A2aR-expressing membrane fraction was obtained by ultra- centrifugation at 100,000 g for 1 hour at 4°C in a Type 70 Ti Fixed-angle rotor (Optima XPN ultracentrifuge, Beckman Coulter Life Sciences). The pellet (membrane fraction) was resuspended at 80 mg/mL wet weight in 50 mM Tris/HCl pH 8, 500 mM NaCl, 10% glycerol. The membranes were homogenized and either used directly for the solubilization of the A2aR or stored at -80°C.

### 1.4. A2AR solubilization using polymers or detergent

Membrane proteins were solubilized from the membrane fraction using the polymers SMA2000, DIBMA free acid, DIBMA monosodium salt, and Glyco-DIBMA or with the detergent DDM, as described previously (Gulamhussein et al., 2020). Membranes from A2aR-expressing cells were resuspended at 40 mg/mL in solubilization buffer (50 mM Tris/HCl pH 8, 500 mM NaCl, 10% glycerol) containing SMA2000, DIBMA free acid, DIBMA monosodium salt or Glyco-DIBMA pH 9.5 at final concentrations of 2.5% (w/v), or the detergent n-Dodecyl β-D-maltoside (DDM) at 2% (w/v). Alternatively, a final concentration of 5% (w/v) of DIBMA polymers was used. The membrane fraction was incubated for 1h at RT with gentle agitation or for 3 h at 4°C for solubilization by detergent. The non-solubilized material was removed by ultra-centrifugation at 100,000 g for 1 h at 4°C to yield supernatants containing A2aR-SMALP, A2aR-DIBMALP or A2aR-DDM.

### 1.5. Purification and analysis of solubilized A2AR preparations

A2aR-SMALP and A2aR-DIBMALP were purified using Ni^2+^-NTA resins essentially as described previously (Jamshad et al., 2015a). The solubilized fractions (10 mL) were incubated overnight with 1 mL of HisPur™ previously equilibrated in 50 mM Tris/HCl pH 8, 500 mM NaCl, 10% glycerol at 4°C. The resin was washed with buffer containing 10 mM or 20 mM imidazole before eluting A2aR-SMALPs with the same buffer containing 60 mM imidazole.

Elution fractions were buffer exchanged into imidazole free buffers using 30 kDa cut-off spin-concentrators. The same procedures were used for A2aR-DDM except that 0.1% DDM was included in the buffers and higher imidazole concentrations were used for elution.

The protein concentrations in the membrane preparation, non-membrane fraction, solubilized fraction, non-solubilized material, and A2aR-SMALP in the elution fractions were quantified using the Pierce^TM^ BCA Protein Assay Kit. Purification was assessed by SDS-PAGE in 12% resolving gels either stained with PageBlue or by transfer onto a PVDF membrane and western blotting with the monoclonal primary antibody anti-Adenosine Receptor A2a (ab79714, Abcam) at 1 µg/mL and the secondary antibody anti-mouse IgG HRP at a 1:3000 dilution. Membranes were imaged with SuperSignal west pico chemiluminescent substrate according to the manufacturer’s instructions (Fisher Scientific, UK).

### 1.6. Analysis of the A2AR protein by LC-MS/MS

After Coomassie staining of the gels obtained by SDS-PAGE, bands of interest corresponding to the A2AR monomer, A2AR dimer, and P. pastoris main contaminants were excised and subjected to in-gel digestion with trypsin or chymotrypsin essentially as described previously (Sousa et al., 2019). Peptides were separated and analysed using an ACQUITY UPLC M-Class LC System (Waters, US) coupled to a 5600 TripleTOF (ABSciex, Warrington, UK) controlled by Analyst software (TF1.5.1, ABSciex, Warrington,UK). The peptide solution (5 µL) was loaded onto a nanoEase MZ Symmetry C18 Trap (180 µm x 20 mm) (Waters, UK), and washed at 20 µl/min for 4 minutes, before separation on a nanoEase MZ Peptide C18 column (15 cm x 75 µm) (Waters, UK) at 35°C. The samples were eluted at 500 nL/min using a gradient elution running from 2% to 45% HPLC Solvent B (99% acetonitrile, and 0.1% formic acid) over 45 min. The peptides were ionized through electrospray ionization (ESI) with a spray voltage of 2.4 kV, source temperature of 150°C, declustering potential of 50 V, and curtain gas setting of 15.

Survey scans were collected in the positive mode from 350 to 2000 Da using a high-resolution TOF-MS mode (MS/MS IDA settings) with 10 ions selected, +2 to +5 charges, dynamic exclusion times of 12s, and rolling collision energy. The data was analysed using the Mascot® search engine (Matrix Science, London, version 2.4.0) to interrogate SwissProt 2021_03 (565,254 sequences; 203,850,821 residues) - Homo sapiens (human) (20,387 sequences) and Other fungi (22,282 sequences).

### 1.7. Analysis of conformational changes by tryptophan fluorescence measurements

Fluorescence measurements were made using a PTI QuantaMaster 300 fluorimeter with continuous Xe arc excitation at 280 nm and emission spectra measured between 290 and 500 nm, essentially as described by (Routledge et al., 2020). The spectra were integrated to obtain the overall intensity. A2aR-SMALP or L-tryptophan as a control (Sigma-Aldrich) were prepared at 50 µg/mL in 50 mM Tris/HCl pH 8, 500 mM NaCl, 10% glycerol containing SMA at 2.5% (w/v). Alternatively, the A2aR-SMALP was treated with acrolein (ACR) or 2,2’-azobis(2- amidinopropane) dihydrochloride (AAPH) at final concentrations of 20 µM, 40 µM, and 100 µM for 24 hours, prior to fluorescence measurements.

To determine the effect of ligand binding on the A2aR, the antagonist ZM241385 or the A2aR agonist NECA were titrated in at concentrations from 1 pM and 100 pM, up to final concentrations of 10 µM and 1 mM, respectively. The resulting intensity data were adjusted to take into account the effect A2aR-SMALP dilution by ligand addition, which was determined by titrating a L-tryptophan control solution with membrane buffer.

The data were analysed using SpectraGryph 1.2 - spectroscopy software and the percentage of integrated tryptophan intensity was calculated taking into account the dilution effect by titration and normalizing to apo A2aR-SMALP or Trp control as 100% of the signal. Data were plotted using GraphPad Prism 8 and expressed as the mean ± standard deviation (SD), for n=3. For statistical data analysis, the percentage of Trp fluorescence at each ligand concentration was compared between the A2aR-SMALP samples treated with different ACR and AAPH concentrations through an ordinary One-way ANOVA and Tukey’s multiple comparisons test. Statistical significance was set at p <0.05.

### 1.8. Thermal unfolding analysis by CPM fluorescence

A2aR-SMALP samples were prepared at final concentration of 150 µg/mL in SMA buffer (50 mM Tris/HCl pH 8, 500 mM NaCl, 10% glycerol, 2% w/v SMA2000); 45 µL of the sample was mixed with 5 µL of 200 µg/mL 7-Diethylamino-3-(4’-Maleimidylphenyl)-4-Methylcoumarin (CPM) was prepared in the same buffer. To study the effect of ligand binding on the thermal stability of the A2aR-SMALP, 1 µM ZM241385 and 10 µM NECA were added to the mixture. To study the effect of acrolein and AAPH, A2aR-SMALP was pre-incubated for 20 min at room temperature with CPM before adding treatments at final concentration of 100 µM and incubating for 1 h at RT. All incubations with CPM were carried out in dark conditions. Thermal unfolding analysis was carried out in white-opaque 96-well PCR plates using a LightCycler® 480 System (Roche Diagnostics). The temperature was equilibrated at 20°C and fluorescence was measured at an excitation λ 465 nm and an emission λ 510 nm from 20 to 99°C with a ramp rate of 3.6°C/min. For calculation of the melting temperatures (Tm), blank-subtracted fluorescence was plotted against temperature, and the first derivative of the blank-subtracted fluorescence (Haffke, 2017) was calculated using GraphPad Prism 8.

### 1.9. Mammalian expression of a hA2AR sequence

A construct for expression of a C-terminal truncated hA2aR in mammalian cells was designed containing FLAG tag, deca-histidine tag, Kozak consensus sequence (AT), unstructured flexible linker (GGSGSG), EcoRI and NotI restriction sites, and was purchased from Eurofins Scientific cloned into a pEX-K248 standard vector (**Supplementary** Figure 4). The C-terminal truncated sequence of the A2aR encodes a stable, functional, and degradation-resistant protein (Prosser et al., 2017). It is well established that the long C-terminal tail is dispensable for receptor folding and dimerization (Hinz et al., 2018) and many A2aR structural studies have used C-terminal truncated sequences (Dore et al., 2011, Garcia-Nafria et al., 2018, Hino et al., 2012, Jaakola et al., 2010).

Following expansion of pcDNA3 empty vector and pEX-K248-hA2aR in competent DH5α *E. coli*, plasmid extraction by miniprep, and digestion of both with EcoRI and NotI, the empty vector and the insert were ligated using T4 DNA ligase (Promega) and transformed DH5a *E. coli* clones containing pcDNA3- hA2aR were selected on plates containing ampicillin using standard protocols.

Human Embryonic Kidney 293T (HEK293T) cells were cultured in DMEM containing 4 mM L-glutamine, 4500 mg/L glucose, 1 mM sodium pyruvate, 1500 mg/L sodium bicarbonate, and phenol red supplemented with 10% foetal bovine serum, penicillin (100 U/ml), and streptomycin (100 µg/ml) at 37°C and 5% CO_2_. HEK293T cells in 6-well plates (9×10^5^ cells / well) were transfected with 100 µL Gibco™ Opti-MEM™ medium mixed with 2 µg of plasmid, 8µL of polyethylenimine (PEI; 1mg/ml, in DPBS pH 7.5), according to the manufacturer’s instructions. The plates were incubated for 48 h at 37°C and 5% CO_2_.

### 1.10. MTT assay for cell viability

HEK293T cells were seeded in 96-well plates at 40000 cells/well in a volume of 100 µL and incubated for 24 h at 37°C and 5% CO_2_ before treatment with final concentrations of 5, 10, 20, 50, 100, 200, and 400 µM acrolein or 0.25, 0.5, 1, 2, 5, 10, and 15 mM AAPH, and incubated for a further 2 or 24 h. The wells were washed once with PBS, resuspended in 90 µL of DMEM without phenol red containing FBS, penicillin and streptomycin. Ten µL of 0.5 mg/mL 3-(4,5- Dimethyl-2-thiazolyl)-2,5-diphenyl-2H-tetrazolium bromide (MTT) solution was added and the plate incubated for 3 h at 37°C and 5% CO^2^, before addition of 100 µL of solubilization solution (10% SDS w/v in 10 mM HCl) to each well. Once formazan crystals were solubilized, the absorbance was measured at 570 nm with a reference wavelength of 670 nm using a Multiskan™ GO Microplate reader.

### 1.11. Cyclic AMP (cAMP) measurements

Cyclic AMP (cAMP) measurements were performed using the AlphaScreen cAMP Detection Kit (PerkinElmer) (Thakur et al., 2023). HEK293T cells (30000 /well) were seeded in 96-well plates and incubated overnight at 37°C and 5% CO2 in 100 µL/well of complete DMEM before transfecting with the pcDNA3-hA2aR plasmid as described above. The cells were treated with vehicle (PBS), 250 µM AAPH, or 20 µM acrolein for 2 or 24 h at 37°C and 5% CO_2_. After treatment, the cells were washed once with PBS and starved in 90 µL/well of pre-warmed and sterile Stimulation Buffer (0.1% w/v BSA, 1 mM 3-isobutyl-1-methylxanthine (IBMX) in phenol-free DMEM) for 1 h at 37°C. The cells were stimulated with 10 µL of 10X ligand half-log serial dilutions for 30 min; the final ligand concentration in the assay was 100 μM to 1 nM for Forskolin and NECA. The ligand-containing medium was aspirated, and 50 μL of ice-cold 98% ethanol was added to each well and allowed to evaporate overnight. Lysis Buffer (75 μL; 0.1% w/v BSA, 0.3% v/v Tween-20, 5 mM HEPES buffer, pH 7.5) was added to each well and incubated with shaking at RT for 10 min. For accurate cAMP measurements, the samples were further diluted ten times in Lysis Buffer. The Biotin-cAMP Acceptor and Streptavidin Donor bead mixes were prepared following the manufacturer’s instructions and incubated for 30 min in the dark before being added to the assay plate. A cAMP standard curve with half-log dilutions from 1 µM to 10 pM in Stimulation Buffer was prepared fresh for each assay. cAMP standards or samples (5 µL) were transferred to a 384-well white opaque Optiplate and, under reduced light conditions, 5 µL of the Acceptor bead and 15 µL of the Streptavidin Donor bead mixtures were added to each well and incubated in the dark at room temperature for 1 h. Fluorescence was measured with excitation at 680 nm and emission 520-620 nm.

### 1.12. Cyclic AMP (cAMP) measurements

All data were analysed and graphs prepared using GraphPad Prism 8. In most cases one-way ANOVA and Tukey’s multiple comparisons test were used with statistical significance set at p <0.05. cAMP data analysed with nonlinear regression and the Kruskal-Wallis test (nonparametric ANOVA) was used to determine statistical differences in pEC50 values.

## 2. Results

### 2.1. hA2aR over-expression in Pichia pastoris, extraction and enrichment

A multi-tagged de-glycosylated C-terminal truncated form of the human A2a receptor was stably expressed in *Pichia pastoris* by transfection with the pPICZB-A2aR plasmid (**Supplementary** Figure 1A **& B**); expression was induced by growth in buffered methanol complex medium (**Supplementary** Figure 1C). Plasma membranes were prepared by cell lysis and ultracentrifugation and solubilized using SMA2000 to form SMALPs. Western blotting of the resulting fractions shows that the A2aR was enriched in the membrane preparation and could be detected in both dimeric and monomeric forms, with the latter predominating (**Supplementary** Figure 2A). Although A2aR was present in the non-solubilized pellet, SMA2000 clearly solubilized the receptor and provided clearer bands. Three different types of DIBMA were also tested and showed that only glycol-DIBMA was effective at extracting the A2AR protein, whereas the detergent DDM gave effective extraction (**Supplementary** Figure 2B**&C**).

To isolate the his-tagged A2AR from other membrane proteins, nickel affinity binding to His-Pur resin with sequential elution by increasing imidazole concentrations was used both for nanodisc- and detergent-extracted membrane preparations. It was noticed that Pichia pastoris contained an endogenous protein that also bound to His-Pur resin and eluted with imidazole. Hence the affinity purification elution steps were optimized for A2aR-SMALPs to separate this contaminant. Good recovery of the A2aR was obtained with 60 mM imidazole without significant contamination, whereas both proteins eluted strongly at higher imidazole concentration (**Figure 1A**). Multiple elution steps at 60 mM imidazole separated the contaminant effectively (**Figure 1B**), and this approach was effective with larger scale SMA-2000 solubilization following lysis with glass beads, whereas glycol-DIBMA solubilization was unsuccessful (data not shown). Purification of A2aR-DDM required higher imidazole levels (150-200 mM) to elute the A2AR and contaminant co-elution could not be prevented (**Figure 1C**). Ultimately, membrane solubilization with SMA2000 enabled the best isolation of the A2aR (**Figure 1D**).

**Figure 1.**
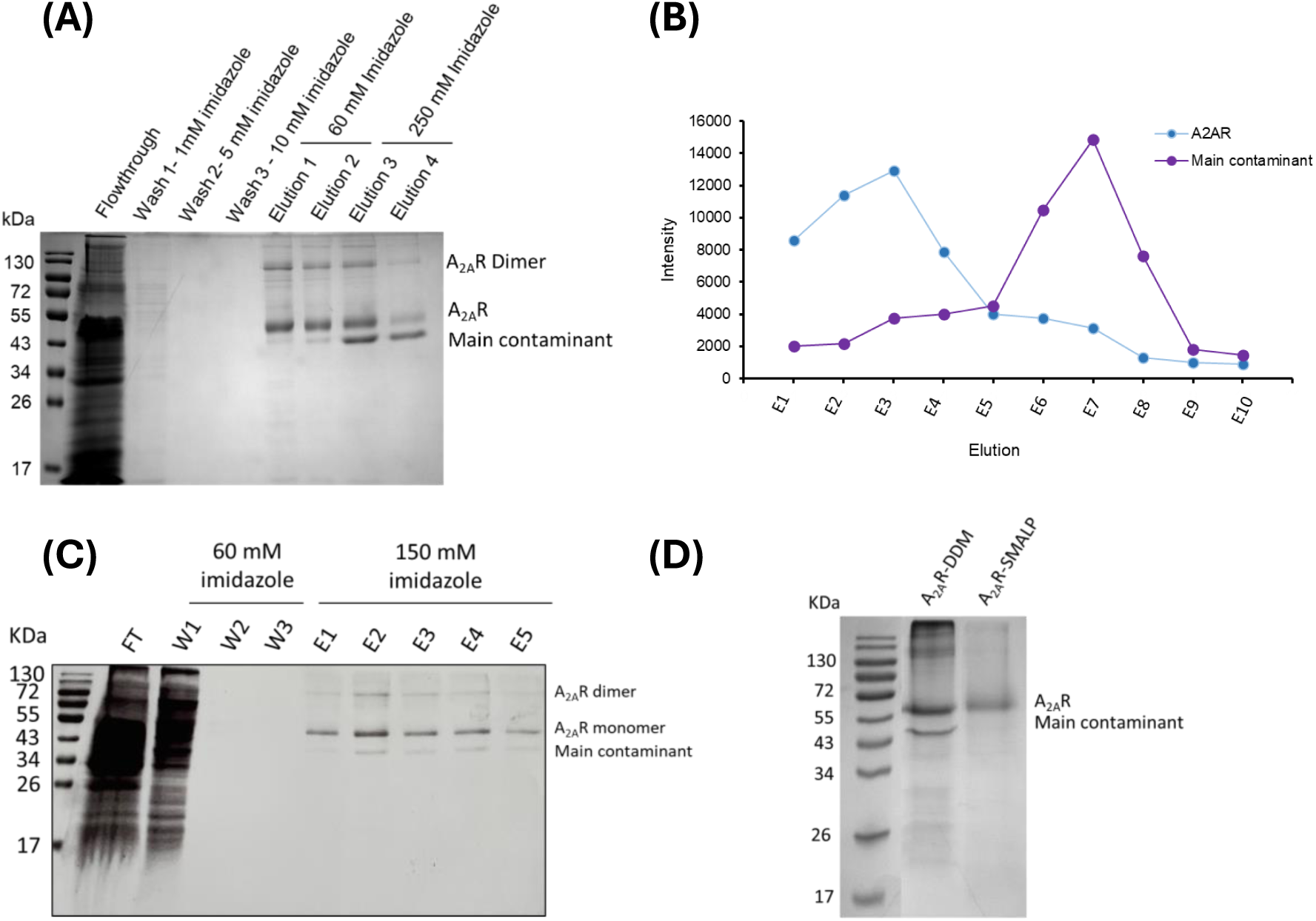
Optimization of A2aR purification from Pichia pastoris using His-Pur Ni-NTA resin. SDS-PAGE with Coomassie staining to visualize the proteins. (A) The SMA-2000 solubilized membrane preparation was loaded onto the His-Pur column, washed with low imidazole concentrations and eluted with 2 aliquots each of 60 and 250 mM imidazole. (B) A2aR-SMALP and contaminant separation by elution using multiple aliquots of 60 mM imidazole; densitometry of the gel using ImageJ. (C) Imidazole elution of A2aR and contaminant after membrane solubilization with DDM. (D) Comparison of the bands corresponding to the A2aR and P. pastoris main contaminant following solubilization of membranes with SMA2000 or DDM and subsequent buffer exchange.

The isolation of the A2aR was confirmed by in-gel digestion of the bands indicated as A2aR dimer and A2aR monomer, and the contaminant observed in the gels was identified as alcohol dehydrogenase 2 (**Table 1**). However, the sequence coverage of the protein was very low; 4% for the A2aR monomer band digested with trypsin (two unique peptides detected), 16% for A2aR monomer band digested with chymotrypsin (seven unique peptides) and 6% for A2aR dimer band digested with chymotrypsin (three unique peptides) (**Supplementary Table 1**). This presumably reflects the challenges of proteomic analysis of membrane proteins.

**Table 1.**
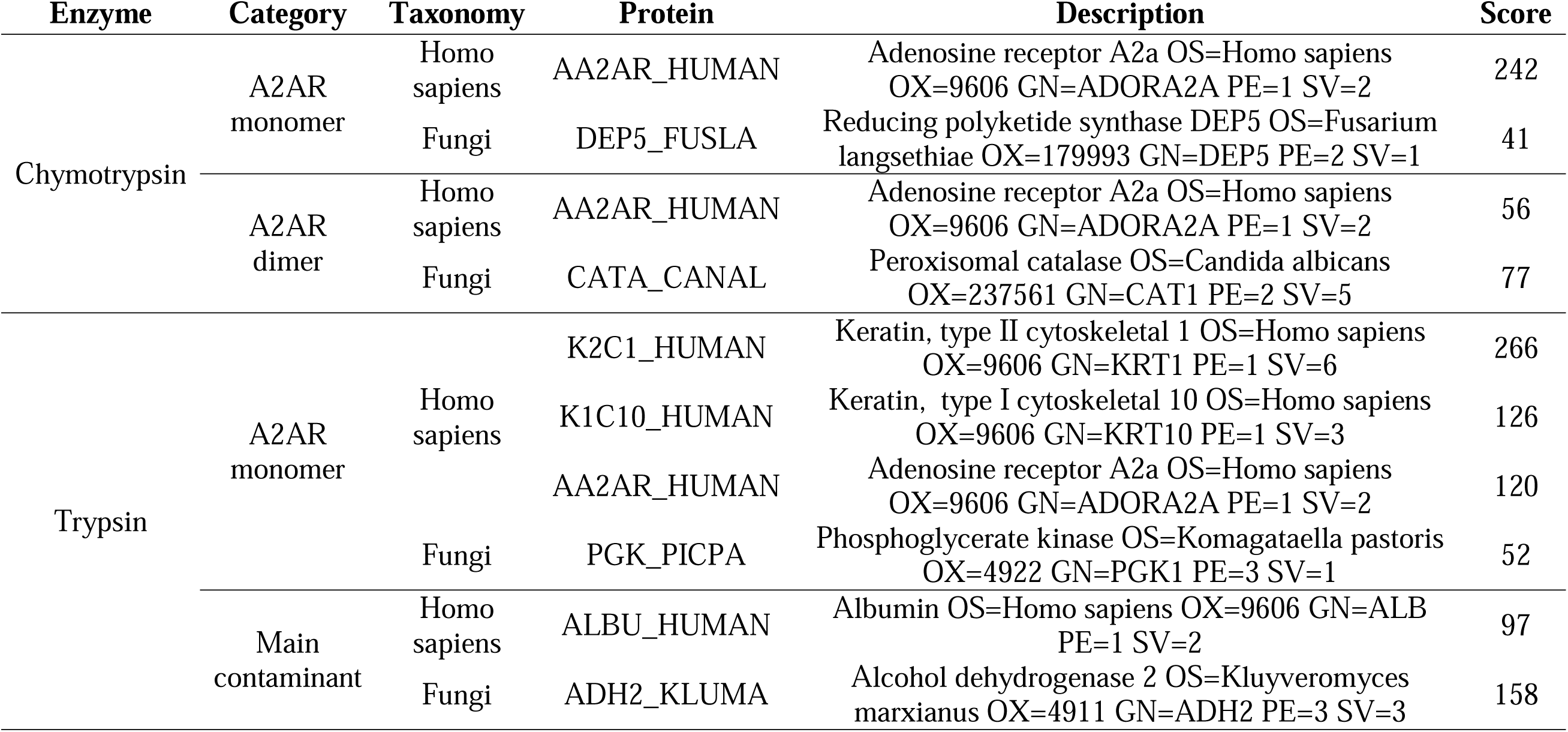
Top proteins identified by LC-MS/MS in the A2aR gel bands from coomassie-stained gels of A2aR- SMALP purified from *Pichia pastoris*.

| Enzyme | Category | Taxonomy | Protein | Description | Score |
| --- | --- | --- | --- | --- | --- |
| Chymotrypsin | A2AR monomer | Homo sapiens | AA2AR_HUMAN | Adenosine receptor A2a OS=Homo sapiens<br>OX=9606 GN=ADORA2A PE=1 SV=2 | 242 |
|  |  | Fungi | DEP5_FUSLA | Reducing polyketide synthase DEP5 OS=Fusarium<br>langsethiae OX=179993 GN=DEP5 PE=2 SV=1 | 41 |
|  | A2AR dimer | Homo sapiens | AA2AR_HUMAN | Adenosine receptor A2a OS=Homo sapiens<br>OX=9606 GN=ADORA2A PE=1 SV=2 | 56 |
|  |  | Fungi | CATA_CANAL | Peroxisomal catalase OS=Candida albicans<br>OX=237561 GN=CAT1 PE=2 SV=5 | 77 |
| Trypsin | A2AR monomer | Homo sapiens | K2C1_HUMAN | Keratin, type II cytoskeletal 1 OS=Homo sapiens<br>OX=9606 GN=KRT1 PE=1 SV=6 | 266 |
|  |  |  | K1C10_HUMAN | Keratin, type I cytoskeletal 10 OS=Homo sapiens<br>OX=9606 GN=KRT10 PE=1 SV=3 | 126 |
|  |  |  | AA2AR_HUMAN | Adenosine receptor A2a OS=Homo sapiens<br>OX=9606 GN=ADORA2A PE=1 SV=2 | 120 |
|  |  | Fungi | PGK_PICPA | Phosphoglycerate kinase OS=Komagataella pastoris<br>OX=4922 GN=PGK1 PE=3 SV=1 | 52 |
|  | Main contaminant | Homo sapiens | ALBU_HUMAN | Albumin OS=Homo sapiens OX=9606 GN=ALB<br>PE=1 SV=2 | 97 |
|  |  | Fungi | ADH2_KLUMA | Alcohol dehydrogenase 2 OS=Kluyveromyces<br>marxianus OX=4911 GN=ADH2 PE=3 SV=3 | 158 |

### 2.2. hA2AR-SMALP was able to undergo conformational changes in response to ligands

In view of SMA-2000 enabling the best isolation of the A2aR, subsequent experiments on A2aR function were carried out using A2aR-SMALPs or A2aR-DDM. Tryptophan fluorescence measurements were performed to determine the effect of ligand binding on the conformational changes in the A2aR-SMALP. In the absence of any ligand, the A2aR-SMALP exhibited a broad fluorescence peak with maximum intensity at ∼340 nm, demonstrating that the fluorescence of tryptophan residues of the A2aR could be measured in low-concentration protein preparations (∼50 µg/mL). An increase in fluorescence intensity, accompanied by a red shift, was observed upon the addition of the antagonist ZM241385 at 1 µM, indicated that the A2aR was folded in the SMALP, and its conformation could be modified upon antagonist binding (**Figure 2A**). In contrast, addition of the agonist NECA at 10 and 100 µM caused decreases in the fluorescence intensity but no emission shift was observed (**Figure 2B**).

**Figure 2.**
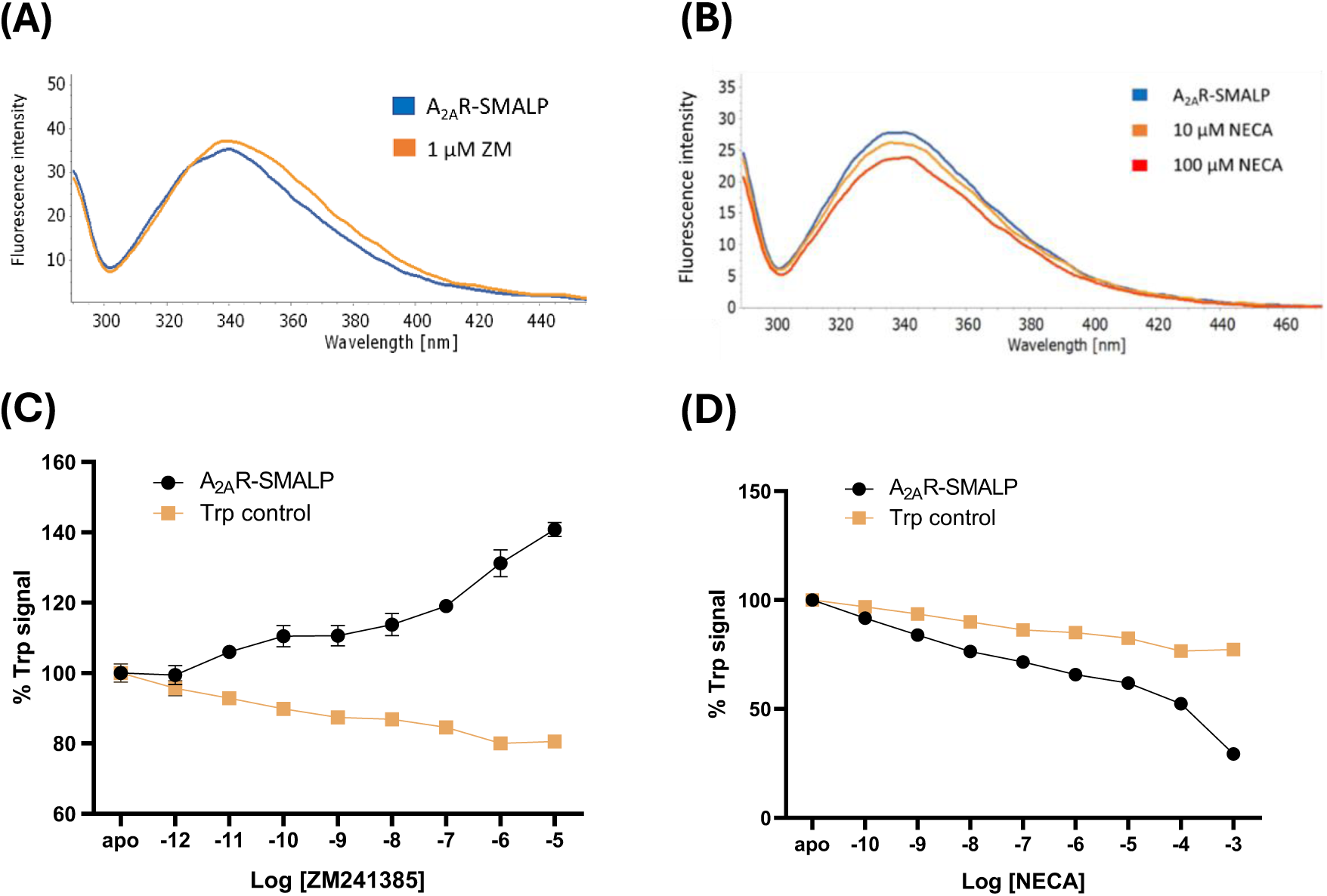
Ligand-induced changes in overall A2aR-SMALP fluorescence. The fluorescence emission of A2aR-SMALP was measured between 290 and 500 nm after excitation at 280 nm before (apo) and after the addition of (A) the antagonist ZM241385 and (B) the agonist NECA (exemplar data from n=3. The integrated fluorescence emission of apo A2aR-SMALP and Trp control after the addition of increasing concentrations of the (C) antagonist ZM241385 and (D) agonist NECA by titration were measured. Results are presented as percentage Trp signal relative to apo A2aR-SMALP or initial Trp control as 100% and taking into account the dilution effect. Data are presented as the mean ± SD (n=3).

To investigate the dose-dependence further, titrations with antagonist and agonist were carried out and the percentage Trp fluorescence signal was calculated by normalizing the fluorescence intensity to the apo-A2aR-SMALP. An increase in the Trp fluorescence signal after the addition of the antagonist to A2aR-SMALP was observed, with greatest effect above 100 nM leading to a 44% final increase in intentsity (**Figure 2C**). ZM241385 addition to the Trp solution control caused a slight decrease in the fluorescent signal, suggesting an effect of the ligand or buffer on tryptophan in solution. Similarly, the agonist NECA progressively decreased the intensity in A2aR-SMALP with the largest effect above 10 µM, showing a greater response than the Trp solution control (**Figure 2D**). Thus, the addition of the agonist NECA to the A2aR-SMALP preparation induced the opposite effect on the Trp fluorescence compared to the antagonist ZM241385. These results suggest the ability of the A2AR with a SMALP to undergo conformational changes from the active to inactive form, and vice versa.

Interestingly, A2aR prepared by solubilizing the membrane with DDM did not show the progressive increase in fluorescence emission in response to ZM241385 titration observed for the A2AR-SMALP and neither did NECA cause a progressive decrease in fluorescence intensity compared to the Trp control in DDM buffer (**Supplementary** Figure 3). This suggested that A2AR-DDM preparation was not able to undergo conformational changes, hence further studies were carried out only with A2AR-SMALP.

### 2.3. hA2AR-SMALP conformational changes were not altered by oxidative treatments

We hypothesized that treatment of the A2aR-SMALP with the radical generator AAPH or reactive carbonyl compound acrolein would affect the ligand binding or ability of the A2aR to undergo conformational changes, either through direct action on the protein or by modification of phospholipids in the SMALP. Consequently, A2aR-SMALP preparations were treated with various concentrations of acrolein or AAPH and titrated with increasing concentrations of ZM241385 or NECA. **Figure 3** shows that neither acrolein nor AAPH caused significant changes to the ligand concentration-response curves: the A2aR was still able to undergo conformational change to the inactive conformation in the presence of ZM241385 and to the active conformation on incubation with NECA.

**Figure 3.**
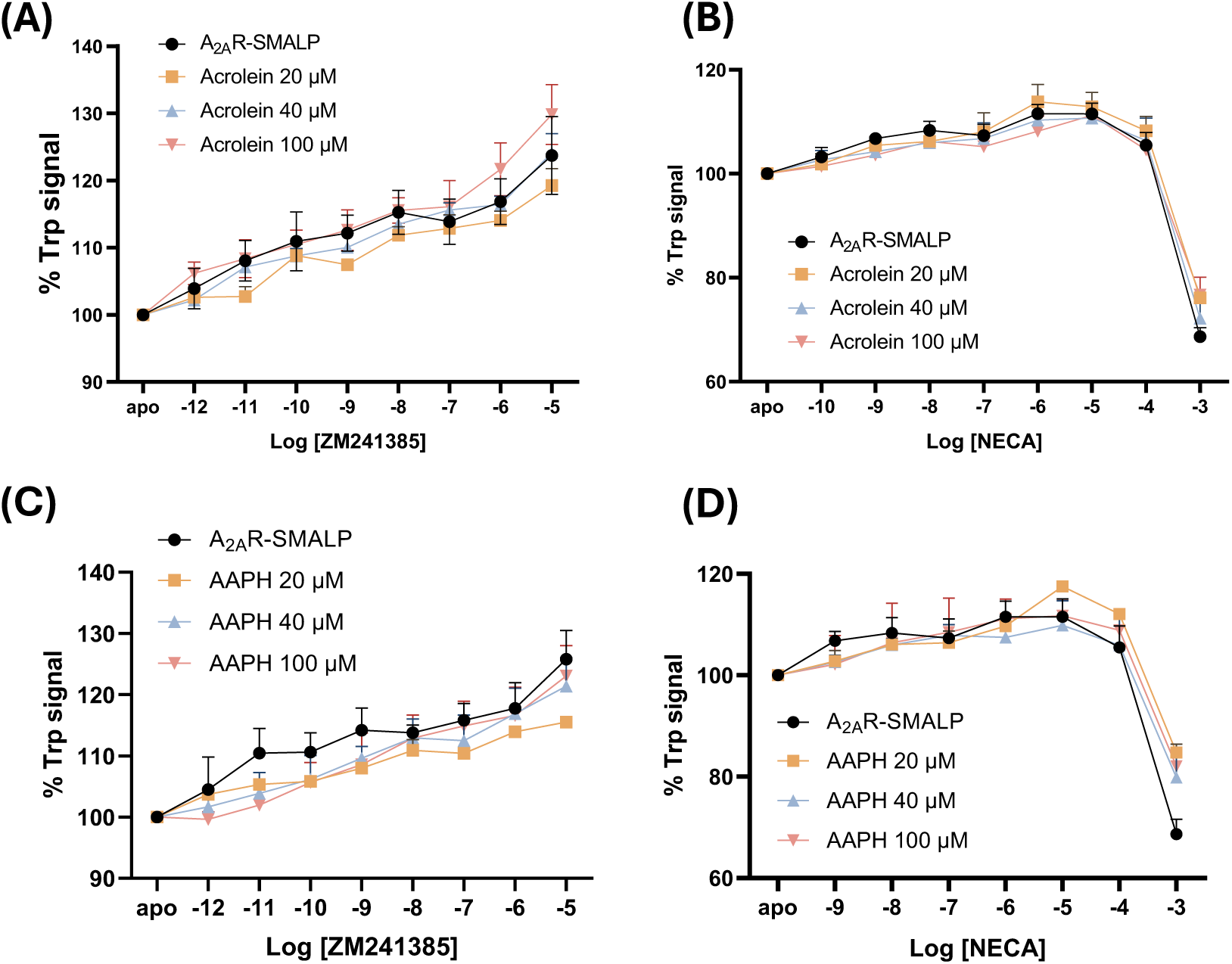
ZM241385- and NECA-induced changes in the overall fluorescence of A2aR- SMALPs treated with acrolein and AAPH. The fluorescence emission of A2AR-SMALP control or subjected to oxidative stress was integrated between 290 and 500 nm after excitation at 280 nm before (apo) and after the addition of increasing concentrations of ligand. (A) Treatment with acrolein and titration with the antagonist ZM241385; (B) treatment with acrolein and titration with the agonist NECA. (C ) Treatment with the radical generator AAPH and titration with the antagonist ZM241385; (D) treatment with AAPH and titration with the agonist NECA. The results are presented as the percentage of Trp signal compared to apo A2AR- SMALP as 100% as described for Figure 2. Statistical differences were determined by one-way ANOVA comparing untreated A2AR-SMALP with AAPH-oxidized receptor (n=3).

### 3.3. Oxidative treatments did not alter thermal unfolding of the hA2aR-SMALP

As no effects on conformational changes of A2AR-SMALP occurred following oxidative stress, the thermostability of the receptor was tested by measuring the fluorescence of the high-affinity probe CPM, which binds to accessible cysteine residues, causing a transition from a non-fluorescent to a fluorescent state and indicating the occurrence of protein unfolding. The apo- and ligand-bound forms of A2AR-SMALP showed a similar pattern of fluorescence increase over the temperature ramp, with unfolding starting at 35-40°C and finishing at ∼60°C (**Figure 4A**). The melting temperatures (T_m_) were determined from the first derivative of the fluorescence; they showed no differences between the 3 samples with all T_m_ values ∼45°C (**Figure 4B**), suggesting that ligand binding did not affect thermal stability. Treatment of the A2AR-SMALP with either 100 µM acrolein or AAPH also had no effect on CPM fluorescence, suggesting that these treatments did not alter the protein stability (Figure 4C&D).

**Figure 4.**
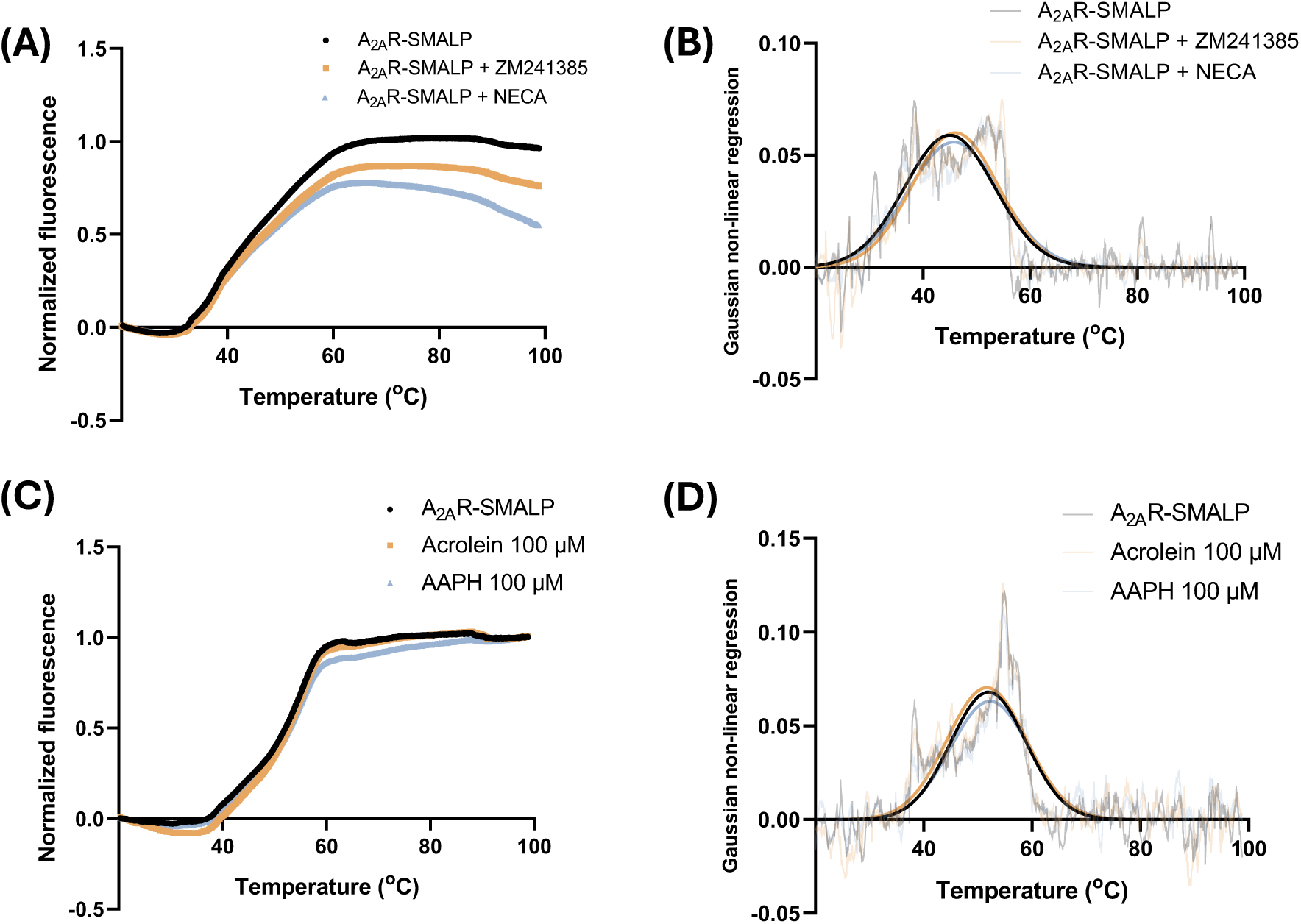
Impact of ligand binding and oxidative treatments on thermal unfolding of the A2aR-SMALP monitored by CPM fluorescence. (A) Fluorescence caused by CPM binding to apo-, ZM241385-bound, and NECA-bound A2AR-SMALP were measured as the temperature increased from 20 to 100oC. (B) The first derivative of fluorescence data from A was plotted against temperature. Gaussian nonlinear regression was performed on the first-derivative data to calculate the Tm. (C) Fluorescence caused by CPM binding to non-treated, ACR-, and AAPH- treated A2AR-SMALP with temperatures increasing from 20 to 100oC. (D) The first derivative of the fluorescence data from C was calculated and plotted against temperature. Gaussian nonlinear regression was performed as above. Combined data from 3 replicates is shown.

### 3.4. Expression of the A2AR in HEK-293 cells

In view of the lack of effect of oxidative treatments on A2aR ligand binding, conformational changes and thermostability in membrane nanodiscs isolated from *Pichia pastoris*, subsequent work focused on downstream signalling in mammalian cells, as yeast do not have the necessary signalling pathways. A construct for mammalian expression was designed and transiently transfected into HEK293 cells, and expression of the A2aR was confirmed by western blotting (**Supplementary** Figure 4) and LC-MS/MS analysis of gel bands digested with trypsin or chymotrypsin (**Supplementary Table 2**).

### 3.5. Selection of oxidative treatment concentrations to maintain HEK-293 cell viability

In order to determine appropriate oxidative treatments that would cause stress without causing loss of viability, MTT assays were carried out. Acrolein decreased cell viability in a concentration-dependent manner after 2 and 24 hours of treatment, with significant decreases at or above 200 µM for 2 hour treatments and 100 µM for 24 hour treatment (**Figure 5A**). AAPH also decreased cell viability in a concentration-dependent manner in cells subjected to 24h treatment with significant effect at 5 mM or higher, whereas it had no significant effect on cells treated for 2h (**Figure 5B**). Consequently, concentrations of 20 µM acrolein and 250 µM AAPH were chosen for all further experiments, to maintain cell viability >85%.

**Figure 5.**
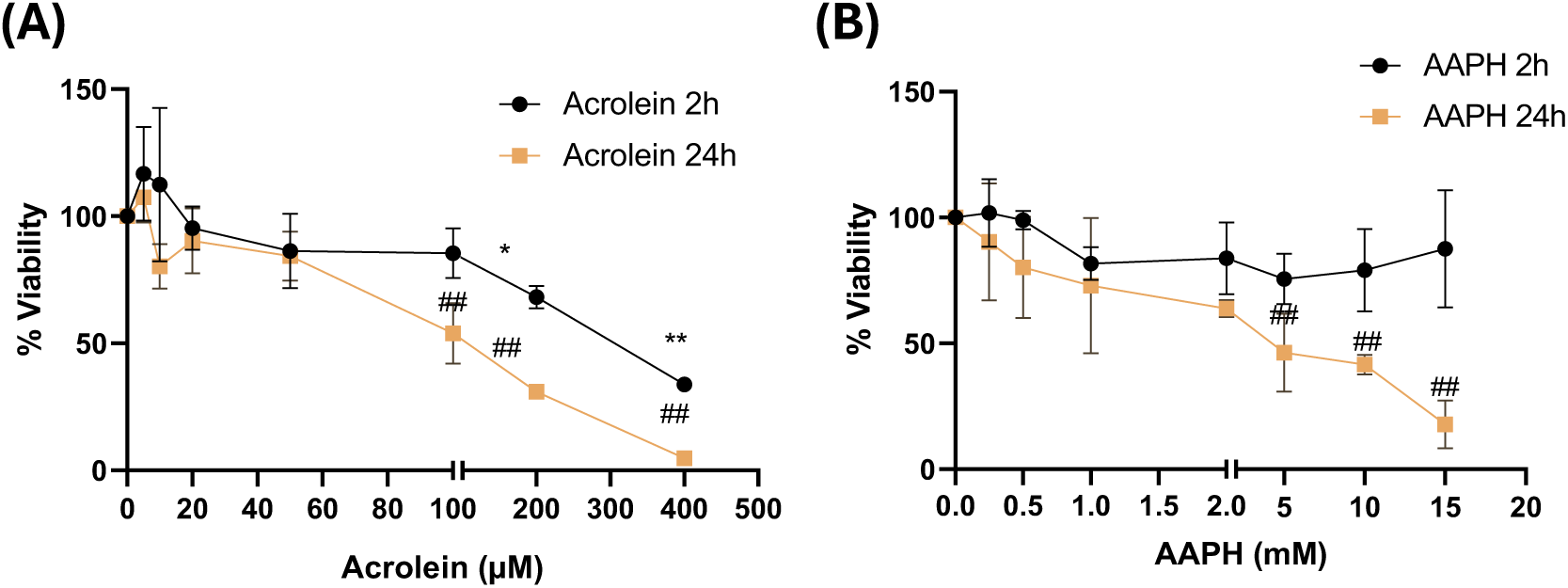
The effect of acrolein and AAPH treatment on HEK293T cell viability. HEK293T cells were seeded at 40000 cells/well onto 96-well plates and treated with increasing concentrations of (A) acrolein or (B) AAPH for 2 and 24 hours. Cell viability was determined by MTT assay after formazan solubilization for 30 minutes at room temperature shaking. Data are presented as mean viability percentage of all replicates (n=3) ± SD. Statistical differences were determined by One-way ANOVA (*p-value<0.05, **p-value<0.01 2h vs control; #p-value<0.05, ##p-value<0.01 24h vs control).

### 3.7. Acrolein and AAPH increase cAMP production in response to NECA

To investigate the effect of oxidative treatments on downstream signalling, cAMP production was measured using the AlphaScreen cAMP detection kit. To confirm that transfection with the A2aR construct enabled signalling and cAMP production, untreated cells were stimulated with NECA as an A2aR agonist (**Figure 6A**). Very low cAMP production was observed with untransfected cells, which could be due to activation of the adenosine A2b receptor, as HEK293T cells natively express this (Gessi et al., 2005) but NECA has a much lower affinity for it (330 nM vs 20 nM for A2aR) (Gao et al., 1999, Matharu et al., 2001). Transfected cells showed more potent NECA-induced production of cAMP (pEC50 6.28±0.34) compared to untransfected cells (pEC50 3.82±0). Direct adenylate cyclase activation by forskolin in untransfected cells resulted in potent cAMP production with a pEC50, as expected (**Figure 6A**). The responses to NECA were then tested in oxidatively stressed cells. Treatments with acrolein and AAPH at both 2 and 24 hours showed significantly higher cAMP production than the untreated controls for all data points, including the apo control in the absence of NECA (**Figure 6B &C**). It was also observed that cAMP in samples treated with oxidants for 24h were 20-fold lower than those in samples treated for 2h, suggesting longer cell incubation times (in the presence or absence of oxidants) caused decreased the responsiveness to the ligand. To investigate whether the increased cAMP production in treated cells could be due to a direct activation of adenylate cyclase, untransfected HEK293T cells were subjected to acrolein and AAPH treatment, and cAMP production was stimulated using forskolin. In contrast to the results obtained after NECA stimulation, forskolin increased cAMP levels in a concentration-dependent manner but oxidative treatments had no effect on cAMP concentration (**Figure 6D &E**). The maximum cAMP values with 100 µM forskolin after both oxidative treatment times were comparable. Oxidative treatments did not cause any changes in forskolin pEC50 values; the NECA pEC50 showed more variability but there were no significant differences (**Table 2**).

**Figure 6.**
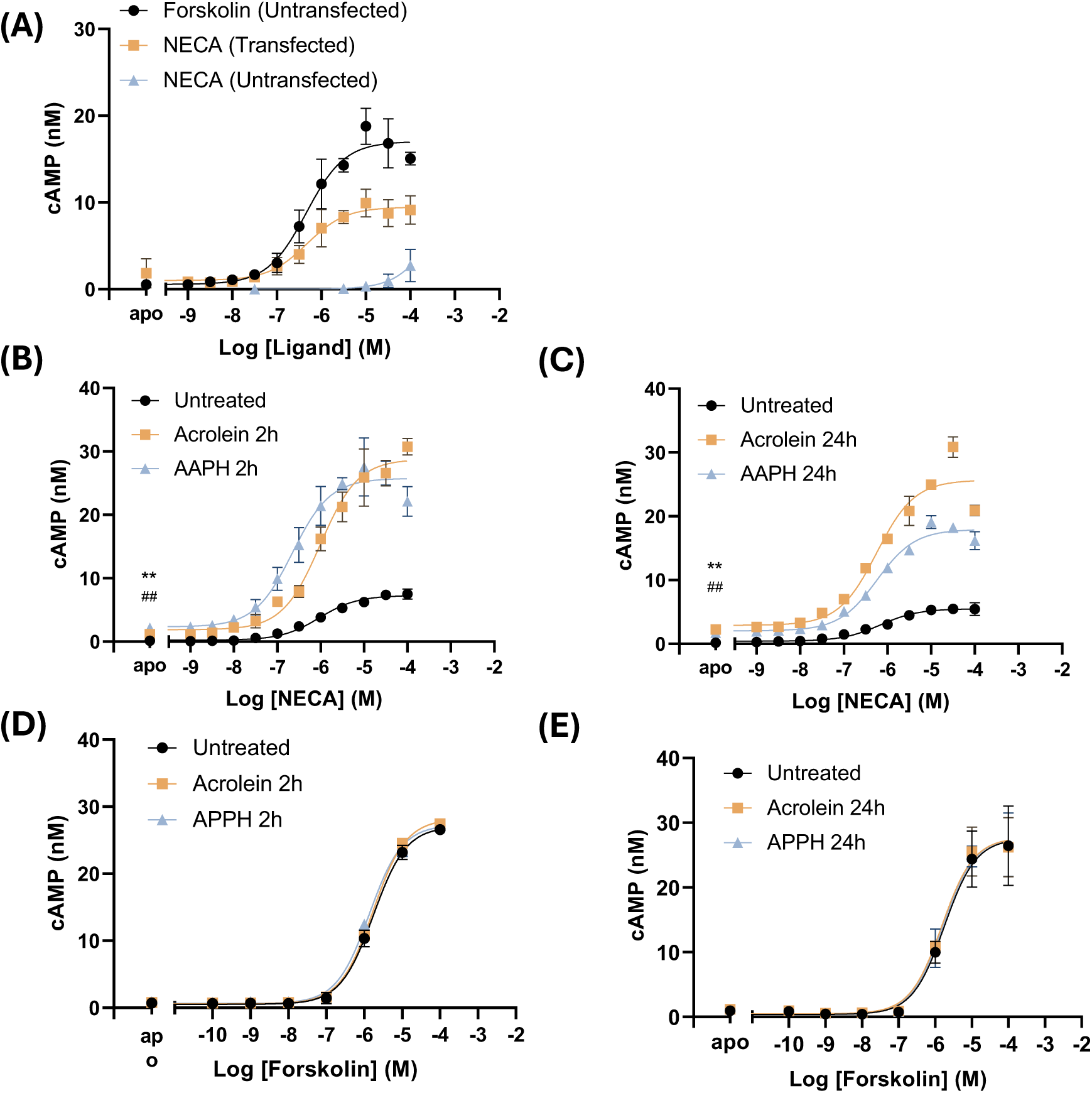
Effect of acrolein and AAPH on cAMP production in HEK293T cells stimulated with NECA or forskolin. The response to forskolin or NECA was compared in untransfected HEK293 cells or cells transiently-transfected with the A2AR plasmid (A) . Transfected HEK293T cells expressing the A2AR were either untreated, treated with acrolein or AAPH for 2 hours (B) or 24 hours (C) and stimulated with NECA. Non-transfected HEK293T cells were either untreated, treated with acrolein or AAPH for 2 hours (D) or 24 hours (E) and stimulated with forskolin. cAMP production was measured as AlphaScreen signal, quantified by comparison with a cAMP standard curve and plotted against the logarithm of NECA concentration (M). Data are presented as the mean of all replicates (n=3) ± SD. Statistical differences were determined by One-way ANOVA followed by Tukey’s multiple comparisons test (*p-value<0.05 acrolein vs untreated).

**Table 2.**
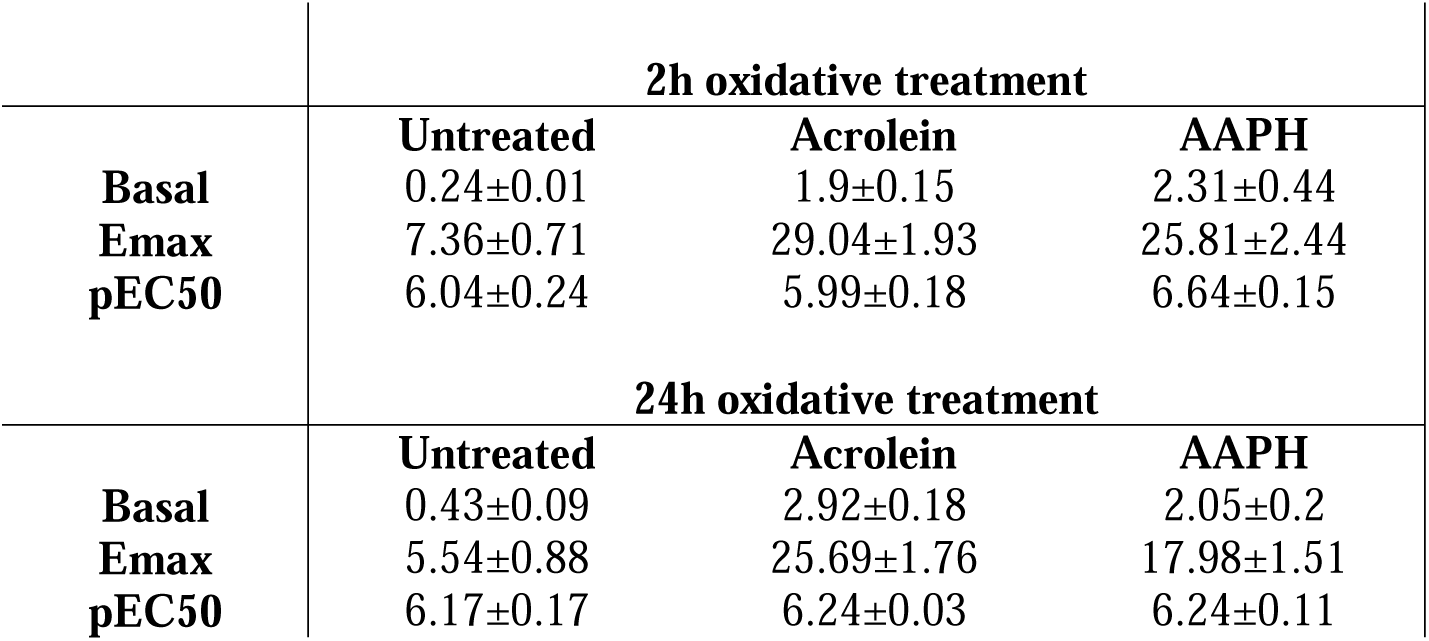
The effect of oxidative treatment on Basal, Emax and pEC50 values, determined from the data in Figure 6.

## Discussion

The overall goal of this study was to investigate the effect of oxidative and lipoxidative stresses on the conformations, activity and signalling of the human adenosine A2a receptor (A2aR). In order to achieve this, the A2aR was over-expressed in two validated expression systems: *P. pastoris* SMD1163 to enable protein purification and HEK293T mammalian cells to study cell signalling. In *P. pastoris*, cell membranes were isolated and solubilized using co-polymers to extract the receptor in nanodics, which retain the native membrane lipid environment. It was found that SMA2000 was the most effective co-polymer for extraction and gave the best purity after extensive optimization of the Ni-NTA affinity purification. Moreover, analysis by tryptophan fluorescence indicated that the A2AR-SMALP was folded and able to undergo ligand-induced conformational changes, whereas A2AR extracted with the detergent DDM, which strips lipids away from the protein, did not show conformational changes. However, oxidative treatments did not alter the ligand-induced conformational changes, nor did they have any effect on the thermostability. In contrast, when HEK293 cells were subjected to sub-lethal oxidative stresses, cAMP levels increased in response to the agonist in a dose-dependent manner that was not dependent on adenylate cyclase activation. This represents the first study to demonstrate effects of oxidative stress on A2AR signalling.

Tryptophan fluorescence provides a convenient approach to study conformational changes in the A2aR and has been used in several previous studies. The C-terminal truncated construct used in this study contained 6 Trp residues, instead of 7 in the native form, but still gave satisfactory fluorescence. The observation that the antagonist ZM241385 caused an increase and red-shift in fluorescence of A2aR-SMALP, corresponding to increased polar environment of the tryptophans, is in agreement a previous report, which suggested that Trp246 (helix 6.48) and Trp268 (helix 7.33) were responsible for the observed increase (Routledge et al., 2020).

Interestingly, they did not observe the decrease in Trp fluorescence upon NECA binding that was found in the current study. Another study reported conformational changes in the tertiary structure of detergent-solubilized A2aR but did not observe changes on antagonist binding, whereas agonist binding caused a blue shift to 326 nm (O’Malley et al., 2010). Our observation that the agonist and antagonist caused opposite changes in the fluorescence emission suggests that the receptor could transit to both active and inactive conformations from its apo state within the SMALP, indicating the existence of mixed populations of apo-encapsulated A2aR-SMALP in different conformational states. This agrees with a previous report that apo-A2aR can be found in different active and inactive states in the absence of ligand (Ye et al., 2016) and the addition of ligands alters the distribution of these conformational states (Prosser et al., 2017). The different results reported by other authors may reflect different populations of the apo-A2aR isolated.

The thermostability measurements confirmed that the A2aR was folded in SMALP, supporting the findings from tryptophan fluorescence. The presence of ligands did not stabilize the protein and result in higher Tm values, in contrast to some previous studies both for SMALPs and detergent extracted A2aR (Haffke, 2017, O’Malley et al., 2010), although other authors have reported stabilizing effects only with certain ligands (Jaakola et al., 2010). Thus, the stabilizing effect of ligands continues to be controversial.

A consistent but surprising finding from the studies of ligand-induced conformational changes, thermostability was that oxidative treatments with acrolein and AAPH had no obvious effects on the equilibrium between conformational states. The original hypothesis was that acrolein would cause direct lipoxidation of the protein while AAPH would either cause direct radical oxidation of the protein or the phospholipids within the SMALP. Phospholipid oxidation can modify the fluidity and lateral pressure within the lipid bilayer, which would be expected to affect A2AR activation; likewise, covalent modification of the protein could interfere with ligand binding in the orthosteric pocket or alter the tertiary structure, affecting conformational changes.

Alternative explanations for the lack of effects are considered below. Either the oxidative modification did not affect the secondary and tertiary structure, allowing the A2AR to be properly folded and active, or the oxidative treatments might not have been sufficient to cause lipid or protein oxidation. However, acrolein concentrations comparable to those used in this study (20 to 100 μM) were sufficient to cause pyruvate kinase lipoxidation, resulting in a significant reduction in protein activity (Sousa et al., 2019), and phosphatase and tensin homolog (PTEN) lipoxidation and inactivation has been reported with 10 μM acrolein (Covey et al., 2010). Previous cell-based studies have observed lipid peroxidation induced by identical AAPH concentrations (Duan et al., 2016, Parathodi Illam et al., 2019). On the other hand, SMA encapsulation may protect both protein and lipid moieties from attack by reactive species, so higher treatment concentration would be needed to cause oxidative damage. A limitation of the current study was that despite digestions with both trypsin and chymotrypsin, very low sequence coverage of the A2aR-SMALP by bottom-up proteomic analysis was obtained, which did not enable mapping of covalent modifications.

In contrast to the lack of effect of oxidative treatments in vitro on detergent- or SMA2000- extracted protein, increased levels of cAMP in response to the agonist NECA were observed in HEK293 cells over-expressing the A2AR and subjected to non-lethal concentrations of acrolein and AAPH. The measurement of cAMP to investigate A2aR function has been well-validated in previous studies (Franco et al., 2019, Igonet et al., 2018, Massink et al., 2015). In the absence of ligands, the increase in cAMP levels was relatively small; a possible explanation for this could be an increase in extracellular adenosine during oxidative stress leading to adenosine receptor activation (Pasquini et al., 2021). In terms of ligand-dependent effects, the NECA pEC50 values observed in the present study were in good agreement with previous pharmacological data (Welihinda et al., 2016, Yan et al., 2003), although lower values have also been reported (Brandon et al., 2006). Although NECA is not specific for the A2aR, the A1R and A3R couple to the Gi protein, which inhibits adenylate cyclase and decreases cAMP production, and therefore cannot be contributing to the observed effect. The A2aR and A2bR both couple to the Gs protein and therefore activate cAMP production through adenylate cyclase, but as NECA induced very low cAMP production in untransfected cells, which natively express A2bR, this suggested that the cAMP production was likely to be dependent on the A2aR. The possibility that direct activation of adenylate cyclase was responsible was ruled out in this study by experiments with its agonist forskolin, which showed no response to oxidative treatments. This disagrees with a previous report in a different cell type (Parkin-mutant fibroblasts) that oxidative stress could cause an increase in basal cAMP levels in via Ca-dependent activation of adenylate cyclase (Tanzarella et al., 2019) but oxidative stress has also been reported to decrease cAMP levels and PKA activity in astrocytes (Shim et al., 2018). Thus, the effect of redox stress on adenylate cyclase remains unclear, possibly owing to different types of oxidants or pathophysiological conditions.

The mechanism by which oxidative treatments lead to NECA-dependent increases in cAMP level is currently unclear. It is known that lipoxidation of proteins can cause a variety conformational changes and alter protein-protein interactions; moreover, certain enzymes can be activated by lipoxidation. Notable examples include the activation of H-ras signalling (Oliva et al., 2003, Renedo et al., 2007), PPARγ (Itoh et al., 2008, Shiraki et al., 2005) and the membrane protein EGFR (Liu et al., 1999) and TRPA (Takahashi et al., 2008)) by HNE or cyclopentenone- containing prostaglandins such as PGA_1_ and 15d-PGJ_2_. It is known that GPCRs can be modified post-translationally by myristoylation, palmitoylation, and isoprenylation (Zhang et al., 2022), and this lipidation can regulate membrane binding, protein trafficking, and the subsequent activation of signalling pathways (Adams et al., 2011). The A2aR can exhibit activity in the absence of agonists, owing to allosteric modulators that shift the equilibrium between states.

Therefore, it is possible that lipoxidation of the A2AR can mimic the effects of lipidation, leading to its activation. A limitation of the study is that the sequence coverage of the extracted A2AR protein was extremely low, as discussed for the experiments on A2AR-SMALP, and likewise it was not possible to detect covalent modifications of the protein to confirm that protein modification had occurred. Alternatively, oxidative stress might alter the membrane lipid profile, which is also known to affect GPCR activity. For example, oxidation of the headgroup of phosphatidylserine can form more negatively charged glycerophosphoacetic acid derivatives, which have been identified in mammalian cells treated with AAPH (Maciel et al., 2014), and might be expected to have increased activation effects on the A2AR.

Although the cellular and *in vitro* results initially appear to be at odds, they are consistent in the sense that oxidative treatments did not adversely affect A2AR-SMALP conformation or stability, while the study of the A2AR in the cellular environment under oxidative stress revealed no impairment in its downstream function. On the other hand, if stimulation of A2AR signalling were due to an altered conformational equilibrium of the protein, it might be expected that this would manifest in altered pEC50 values *in vivo* and in a differential response to agonist and antagonist titrations *in vitro*. However, although widely used, tryptophan fluorescence measurements are not the most sensitive measures of conformational states, so ultimately sensitive and structurally informative methods would be required to confirm the molecular mechanisms.

## Conclusion

While the effect of activating or inhibiting the A2AR on inflammatory conditions and its potential for therapy has been extensively, very few studies have directly investigated the effect of oxidative stress or lipoxidation on the A2AR. We believe this is the first report of the impact of oxidative stress on its conformational changes and downstream signalling. The upregulation of A2AR signalling in response to oxidative treatments implies the potential to modulate oxidative stress and inflammation and could represent a negative feedback loop to control inflammation (Castro et al., 2020, Tavares et al., 2020). It also illustrates the concept of beneficial effects of oxidation at appropriate concentrations and the dichotomies that can occur in redox signalling.

## Acknowledgements

We gratefully acknowledge the kind gift of pPICZB containing the de-glycosylated C-terminal truncated A2aR from Prof Roslyn Bill, Aston Institute for Membrane Excellence. We thank Dr Alice Rothnie for the kind gift of SMA2000 and Dr Lucas Unger for support with the thermal unfolding experiments.

## Author Contributions

CMS, JS, ARP and DRP coordinated the study. ICM carried out most of the practical work. JG carried out some of the cAMP measurements. CMS, ICM, ARP and DRP contributed to manuscript writing; all authors have read the final version of the manuscript.

## Abbreviations

AAPH: 2,2’-azobis(2-amidinopropane) dihydrochloride
A2aR: adenosine A2a receptor
ACR: acrolein
DDM: n-Dodecyl β-D-maltoside
DIBMA: diisobutylene maleic acid
GPCRs: G- protein coupled receptors
NECA: 5′-(N-Ethylcarboxamido) adenosine
SMALP: styrene maleic acid lipid particle
Trp: tryptophan
TM: transmembrane helixes.

## References

1. Adams, M. N., Christensen, M. E., He, Y., Waterhouse, N. J. & Hooper, J. D. 2011. The role of palmitoylation in signalling, cellular trafficking and plasma membrane localization of protease-activated receptor-2. PLoS One, 6, e28018.

2. Aldini, G., Domingues, M. R., Spickett, C. M., Domingues, P., Altomare, A., Sánchez-Gómez, F. J., Oeste, C. L. & Pérez-Sala, D. 2015. Protein lipoxidation: Detection strategies and challenges. Redox Biology, 5, 253–266.

3. Baccouch, R., Rascol, E., Stoklosa, K. & Alves, I. D. 2022. The role of the lipid environment in the activity of G protein coupled receptors. Biophys Chem, 285, 106794.

4. Brandon, C. I., Vandenplas, M., Dookwah, H. & Murray, T. F. 2006. Cloning and pharmacological characterization of the equine adenosine A3 receptor. J Vet Pharmacol Ther, 29, 255–63.

5. Bruzzese, A., Dalton, J. A. R. & Giraldo, J. 2020. Insights into adenosine A2A receptor activation through cooperative modulation of agonist and allosteric lipid interactions. PLoS Comput Biol, 16, e1007818.

6. Burstein, E. A., Vedenkina, N. S. & Ivkova, M. N. 1973. Fluorescence and the location of tryptophan residues in protein molecules. Photochem Photobiol, 18, 263–79.

7. Castro, C. M., Corciulo, C., Solesio, M. E., Liang, F., Pavlov, E. V. & Cronstein, B. N. 2020. Adenosine A2A receptor (A2AR) stimulation enhances mitochondrial metabolism and mitigates reactive oxygen species-mediated mitochondrial injury. FASEB J, 34, 5027–5045.

8. Covey, T. M., Edes, K., Coombs, G. S., Virshup, D. M. & Fitzpatrick, F. A. 2010. Alkylation of the tumor suppressor PTEN activates Akt and beta-catenin signaling: a mechanism linking inflammation and oxidative stress with cancer. PLoS One, 5, e13545.

9. Dawaliby, R., Trubbia, C., Delporte, C., Masureel, M., VAN Antwerpen, P., Kobilka, B. K. & Govaerts, C. 2016. Allosteric regulation of G protein-coupled receptor activity by phospholipids. Nat Chem Biol, 12, 35–9.

10. De lera Ruiz, M., Lim, Y. H. & Zheng, J. 2014. Adenosine A2A receptor as a drug discovery target. J Med Chem, 57, 3623–50.

11. Domingues, R. M., Domingues, P., Melo, T., Perez-Sala, D., Reis, A. & Spickett, C. M. 2013. Lipoxidation adducts with peptides and proteins: deleterious modifications or signaling mechanisms? J Proteomics, 92, 110–31.

12. Dore, A. S., Robertson, N., Errey, J. C., Ng, I., Hollenstein, K., Tehan, B., Hurrell, E., Bennett, K., Congreve, M., Magnani, F., Tate, C. G., Weir, M. & Marshall, F. H. 2011. Structure of the adenosine A(2A) receptor in complex with ZM241385 and the xanthines XAC and caffeine. Structure, 19, 1283–93.

13. Duan, W. J., Li, Y. F., Liu, F. L., Deng, J., Wu, Y. P., Yuan, W. L., Tsoi, B., Chen, J. L., Wang, Q., Cai, S. H., Kurihara, H. & He, R. R. 2016. A SIRT3/AMPK/autophagy network orchestrates the protective effects of trans-resveratrol in stressed peritoneal macrophages and RAW 264.7 macrophages. Free Radic Biol Med, 95, 230–42.

14. Egea, J., Fabregat, I., Frapart, Y. M., Ghezzi, P., Görlach, A., Kietzmann, T., Kubaichuk, K., Knaus, U. G., Lopez, M. G., Olaso-Gonzalez, G., Petry, A., Schulz, R., Vina, J., Winyard, P., Abbas, K., Ademowo, O. S., Afonso, C. B., Andreadou, I., Antelmann, H., Antunes, F., Aslan, M., Bachschmid, M. M., Barbosa, R. M., Belousov, V., Berndt, C., Bernlohr, D., Bertrán, E., Bindoli, A., Bottari, S. P., Brito, P. M., Carrara, G., Casas, A. I., Chatzi, A., Chondrogiannni, N., Conrad, M., Cooke, M. S., Costa, J. G., Cuadrado, A., Dang, P. M. C., DE Smet, B., Debelec-Butuner, B., Dias, I. H. K., Dunn, J. D., Edson, A. J., EL ASSAR, M., EL-Benna, J., Ferdinandy, P., Fernandes, A. S., Fladmark, K. E., Förstermann, U., Giniatullin, R., Giricz, Z., Görbe, A., Griffiths, H., Hampl, V., Hanf, A., Herget, J., Hernansanz-Agustín, P., Hillion, M., Huang, J., Ilikay, S., Jansen-Dürr, P., Jaquet, V., Joles, J. A., Kalyanaraman, B., Kaminskyy, D., Karbaschi, M., Kleanthous, M., Klotz, L. O., Korac, B., Korkmaz, K. S., Koziel, R., Kracun, D., Krause, K. H., Kren, V., Krieg, T., Laranjinha, J., Lazou, A., Li, H., Martínez- Ruiz, A., Matsui, R., Mcbean, G. J., Meredith, S. P., Messens, J., Miguel, V., Mikhed, Y., Milisav, I., Milkovic, L., Miranda-Vizuete, A., Mojovic, M., Monsalve, M., Mouthuay, P. A., Mulvey, J., Münzel, T., Muzykantov, V., Nguyen, I. T. N., Oelze, M., Oliveira, N. G., Palmeira, C. M., Papaevgeniou, N., et al. 2018. European contribution to the study of ROS: A summary of the findings and prospects for the future from the COST action BM1203 (EU-ROS) (vol 13, pg 94, 2017). Redox Biology, 14, 694–696.

15. EL-Shamarka, M. E. A., Kozman, M. R. & Messiha, B. A. S. 2020. The protective effect of inosine against rotenone-induced Parkinson’s disease in mice; role of oxidonitrosative stress, ERK phosphorylation, and A2AR expression. Naunyn Schmiedebergs Arch Pharmacol, 393, 1041–1053.

16. Franco, R., Reyes-Resina, I., Aguinaga, D., Lillo, A., Jimenez, J., Raich, I., Borroto-Escuela, D. O., Ferreiro-Vera, C., Canela, E. I., Sanchez de Medina, V., Del ser-Badia, A., Fuxe, K., Saura, C. A. & Navarro, G. 2019. Potentiation of cannabinoid signaling in microglia by adenosine A(2A) receptor antagonists. Glia, 67, 2410–2423.

17. Gao, Z., Chen, T., Weber, M. J. & Linden, J. 1999. A2B adenosine and P2Y2 receptors stimulate mitogen-activated protein kinase in human embryonic kidney-293 cells. cross- talk between cyclic AMP and protein kinase c pathways. J Biol Chem, 274, 5972–80.

18. GARCIA-Nafria, J., Lee, Y., Bai, X., Carpenter, B. & Tate, C. G. 2018. Cryo-EM structure of the adenosine A(2A) receptor coupled to an engineered heterotrimeric G protein. Elife, 7.

19. Geiger, J., Sexton, R., AL-Sahouri, Z., Lee, M. Y., Chun, E., Harikumar, K. G., Miller, L. J., Beckstein, O. & Liu, W. 2021. Evidence that specific interactions play a role in the cholesterol sensitivity of G protein-coupled receptors. Biochim Biophys Acta Biomembr, 1863, 183557.

20. Gessi, S., Varani, K., Merighi, S., Cattabriga, E., Pancaldi, C., Szabadkai, Y., Rizzuto, R., Klotz, K. N., Leung, E., MAC Lennan, S., Baraldi, P. G. & Borea, P.A. 2005. Expression, pharmacological profile, and functional coupling of A2B receptors in a recombinant system and in peripheral blood cells using a novel selective antagonist radioligand, [3H]MRE 2029-F20. Mol Pharmacol, 67, 2137–47.

21. Ghisaidoobe, A. B. & Chung, S. J. 2014. Intrinsic tryptophan fluorescence in the detection and analysis of proteins: a focus on Forster resonance energy transfer techniques. Int J Mol Sci, 15, 22518–38.

22. Goulding, J., May, L. T. & Hill, S. J. 2018. Characterisation of endogenous A(2A) and A(2B) receptor-mediated cyclic AMP responses in HEK 293 cells using the GloSensor biosensor: Evidence for an allosteric mechanism of action for the A(2B)-selective antagonist PSB 603. Biochem Pharmacol, 147, 55–66.

23. Gulamhussein, A. A., Uddin, R., Tighe, B. J., Poyner, D. R. & Rothnie, A. J. 2020. A comparison of SMA (styrene maleic acid) and DIBMA (di-isobutylene maleic acid) for membrane protein purification. Biochim Biophys Acta Biomembr, 1862, 183281.

24. Haffke, M. R., G.; Boivineau, J.; MÜNCH A.; Jaakola, V. . 2017. *Thermal Unfolding of GPCRs NanoDSF : Label-Free Thermal Unfolding Assay of G Protein- Coupled Receptors for Compound Screening and Buffer Composition Optimization* [Online]. Available: https://resources.nanotempertech.com/application-notes/application-note-nt-pr-008-thermal-unfolding-of-gpcrs [Accessed NT-PR-008-01].

25. Hino, T., Arakawa, T., Iwanari, H., Yurugi-Kobayashi, T., Ikeda-Suno, C., Nakada-Nakura, Y., Kusano-Arai, O., Weyand, S., Shimamura, T., Nomura, N., Cameron, A. D., Kobayashi, T., Hamakubo, T., Iwata, S. & Murata, T. 2012. G-protein-coupled receptor inactivation by an allosteric inverse-agonist antibody. Nature, 482, 237–40.

26. Hinz, S., Navarro, G., Borroto-Escuela, D., Seibt, B. F., Ammon, Y. C., DE Filippo, E., Danish, A., Lacher, S. K., Červinková, B., Rafehi, M., Fuxe, K., Schiedel, A. C., Franco, R. & Müller, C.E. 2018. Adenosine A(2A). receptor ligand recognition and signaling is blocked by A(2B) receptors. Oncotarget, 9, 13593–13611.

27. Igonet, S., Raingeval, C., Cecon, E., Pucic-Bakovic, M., Lauc, G., Cala, O., Baranowski, M., Perez, J., Jockers, R., Krimm, I. & Jawhari, A. 2018. Enabling STD-NMR fragment screening using stabilized native GPCR: A case study of adenosine receptor. Sci Rep, 8, 8142.

28. Ikram, M., Park, T. J., Ali, T. & Kim, M. O. 2020. Antioxidant and Neuroprotective Effects of Caffeine against Alzheimer’s and Parkinson’s Disease: Insight into the Role of Nrf-2 and A2AR Signaling. Antioxidants (Basel), 9.

29. Itoh, T., Fairall, L., Amin, K., Inaba, Y., Szanto, A., Balint, B. L., Nagy, L., Yamamoto, K. & Schwabe, J. W. 2008. Structural basis for the activation of PPARgamma by oxidized fatty acids. Nat Struct Mol Biol, 15, 924–31.

30. Jaakola, V. P., Lane, J. R., Lin, J. Y., Katritch, V., Ijzerman, A. P. & Stevens, R. C. 2010. Ligand binding and subtype selectivity of the human A(2A) adenosine receptor: identification and characterization of essential amino acid residues. J Biol Chem, 285, 13032–44.

31. Jamshad, M., Charlton, J., Lin, Y. P., Routledge, S. J., Bawa, Z., Knowles, T. J., Overduin, M., Dekker, N., Dafforn, T. R., Bill, R. M., Poyner, D. R. & Wheatley, M. 2015a. G-protein coupled receptor solubilization and purification for biophysical analysis and functional studies, in the total absence of detergent. Biosci Rep, 35.

32. Jamshad, M., Grimard, V., Idini, I., Knowles, T. J., Dowle, M. R., Schofield, N., Sridhar, P., Lin, Y. P., Finka, R., Wheatley, M., Thomas, O. R., Palmer, R. E., Overduin, M., Govaerts, C., Ruysschaert, J. M., Edler, K. J. & Dafforn, T.R. 2015b. Structural analysis of a nanoparticle containing a lipid bilayer used for detergent-free extraction of membrane proteins. Nano Res, 8, 774–789.

33. Jones, A. J. Y., Gabriel, F., Tandale, A. & Nietlispach, D. 2020. Structure and Dynamics of GPCRs in Lipid Membranes: Physical Principles and Experimental Approaches. Molecules, 25.

34. Knowles, T. J., Finka, R., Smith, C., Lin, Y. P., Dafforn, T. & Overduin, M. 2009. Membrane proteins solubilized intact in lipid containing nanoparticles bounded by styrene maleic acid copolymer. J Am Chem Soc, 131, 7484–5.

35. Lebon, G., Warne, T., Edwards, P. C., Bennett, K., Langmead, C. J., Leslie, A. G. & Tate, C. G. 2011. Agonist-bound adenosine A2A receptor structures reveal common features of GPCR activation. Nature, 474, 521–5.

36. Leone, R. D. & Emens, L. A. 2018. Targeting adenosine for cancer immunotherapy. J Immunother Cancer, 6, 57.

37. Li, Y. F., Ouyang, S. H., Tu, L. F., Wang, X., Yuan, W. L., Wang, G. E., Wu, Y. P., Duan, W. J., Yu, H. M., Fang, Z. Z., Kurihara, H., Zhang, Y. & He, R. R. 2018. Caffeine Protects Skin from Oxidative Stress-Induced Senescence through the Activation of Autophagy. Theranostics, 8, 5713–5730.

38. Liu, W., Akhand, A. A., Kato, M., Yokoyama, I., Miyata, T., Kurokawa, K., Uchida, K. & Nakashima, I. 1999. 4-hydroxynonenal triggers an epidermal growth factor receptor-linked signal pathway for growth inhibition. J Cell Sci, 112 ( Pt 14), 2409–17.

39. Logez, C., Damian, M., Legros, C., Dupre, C., Guery, M., Mary, S., Wagner, R., M’kadmi, C., Nosjean, O., Fould, B., Marie, J., Fehrentz, J. A., Martinez, J., Ferry, G., Boutin, J. A. & Baneres, J. L. 2016. Detergent-free Isolation of Functional G Protein-Coupled Receptors into Nanometric Lipid Particles. Biochemistry, 55, 38–48.

40. Maciel, E., Neves, B. M., Santinha, D., Reis, A., Domingues, P., TERESA Cruz, M., Pitt, A. R., Spickett, C. M. & Domingues, M.R. 2014. Detection of phosphatidylserine with a modified polar head group in human keratinocytes exposed to the radical generator AAPH. Arch Biochem Biophys, 548, 38–45.

41. Manglik, A. & Kruse, A. C. 2017. Structural Basis for G Protein-Coupled Receptor Activation. Biochemistry, 56, 5628–5634.

42. Massink, A., Gutierrez-de-Teran, H., Lenselink, E. B., Ortiz Zacarias, N. V., Xia, L., Heitman, L. H., Katritch, V., Stevens, R. C. & Ap, I. J. 2015. Sodium ion binding pocket mutations and adenosine A2A receptor function. Mol Pharmacol, 87, 305–13.

43. Matharu, A. L., Mundell, S. J., Benovic, J. L. & Kelly, E. 2001. Rapid agonist-induced desensitization and internalization of the A(2B) adenosine receptor is mediated by a serine residue close to the COOH terminus. J Biol Chem, 276, 30199–207.

44. O’malley, M. A., Naranjo, A. N., Lazarova, T. & Robinson, A. S. 2010. Analysis of adenosine A(2)a receptor stability: effects of ligands and disulfide bonds. Biochemistry, 49, 9181–9.

45. Oeste, C. L., Díez-Dacal, B., Bray, F., DE Lacoba, M.G., De la Torre, B. G., Andreu, D., Ruiz-Sánchez, A. J., Pérez-Inestrosa, E., García- Domínguez, C. A., Rojas, J. M. & Pérez-Sala, D. 2011. The C-Terminus of H-Ras as a Target for the Covalent Binding of Reactive Compounds Modulating Ras-Dependent Pathways. Plos One, 6.

46. Oliva, J. L., Perez-Sala, D., Castrillo, A., Martinez, N., Canada, F. J., Bosca, L. & Rojas, J. M. 2003. The cyclopentenone 15-deoxy-delta 12,14- prostaglandin J2 binds to and activates H-Ras. Proc Natl Acad Sci U S A, 100, 4772–7.

47. Paila, Y. D. & Chattopadhyay, A. 2009. The function of G-protein coupled receptors and membrane cholesterol: specific or general interaction? Glycoconj J, 26, 711–20.

48. PARATHODI Illam, S., Hussain, A., Elizabeth, A., Narayanankutty, A. & Raghavamenon, A.C. 2019. Natural combination of phenolic glycosides from fruits resists pro-oxidant insults to colon cells and enhances intrinsic antioxidant status in mice. Toxicol Rep, 6, 703–711.

49. Pasquini, S., Contri, C., Borea, P. A., Vincenzi, F. & Varani, K. 2021. Adenosine and Inflammation: Here, There and Everywhere. Int J Mol Sci, 22.

50. Prosser, R. S., Ye, L., Pandey, A. & Orazietti, A. 2017. Activation processes in ligand-activated G protein-coupled receptors: A case study of the adenosine A(2A) receptor. Bioessays, 39.

51. REAL Hernandez, L. M. & Levental, I. 2023. Lipid packing is disrupted in copolymeric nanodiscs compared with intact membranes. Biophys J, 122, 2256–2266.

52. Reis, A. & Spickett, C. M. 2012. Chemistry of phospholipid oxidation. Biochim Biophys Acta, 1818, 2374–87.

53. Renedo, M., Gayarre, J., Garcia-Dominguez, C. A., Perez-Rodriguez, A., Prieto, A., Canada, F. J., Rojas, J. M. & Perez-Sala, D. 2007. Modification and activation of Ras proteins by electrophilic prostanoids with different structure are site-selective. Biochemistry, 46, 6607–16.

54. Routledge, S. J., Jamshad, M., Little, H. A., Lin, Y. P., Simms, J., Thakker, A., Spickett, C. M., Bill, R. M., Dafforn, T. R., Poyner, D. R. & Wheatley, M. 2020. Ligand-induced conformational changes in a SMALP-encapsulated GPCR. Biochim Biophys Acta Biomembr, 1862, 183235.

55. Shim, M. S., Kim, K. Y., Bu, J. H., Nam, H. S., Jeong, S. W., Park, T. L., Ellisman, M. H., Weinreb, R. N. & Ju, W.K. 2018. Elevated intracellular cAMP exacerbates vulnerability to oxidative stress in optic nerve head astrocytes. Cell Death Dis, 9, 285

56. Shiraki, T., Kamiya, N., Shiki, S., Kodama, T. S., Kakizuka, A. & Jingami, H. 2005. Alpha,beta-unsaturated ketone is a core moiety of natural ligands for covalent binding to peroxisome proliferator-activated receptor gamma. J Biol Chem, 280, 14145–53.

57. Sousa, B. C., Ahmed, T., Dann, W. L., Ashman, J., Guy, A., Durand, T., Pitt, A. R. & Spickett, C. M. 2019. Short-chain lipid peroxidation products form covalent adducts with pyruvate kinase and inhibit its activity in vitro and in breast cancer cells. Free Radic Biol Med, 144, 223–233.

58. Sousa, B. C., Pitt, A. R. & Spickett, C. M. 2017. Chemistry and analysis of HNE and other prominent carbonyl-containing lipid oxidation compounds. Free Radic Biol Med, 111, 294–308.

59. Spickett, C. M. & Pitt, A. R. 2020. Modification of proteins by reactive lipid oxidation products and biochemical effects of lipoxidation. Essays Biochem, 64, 19–31.

60. Steyaert, J. & Kobilka, B. K. 2011. Nanobody stabilization of G protein-coupled receptor conformational states. Curr Opin Struct Biol, 21, 567–72.

61. Takahashi, N., Mizuno, Y., Kozai, D., Yamamoto, S., Kiyonaka, S., Shibata, T., Uchida, K. & Mori, Y. 2008. Molecular characterization of TRPA1 channel activation by cysteine-reactive inflammatory mediators. Channels (Austin), 2, 287–98.

62. Tanzarella, P., Ferretta, A., Barile, S. N., Ancona, M., DE Rasmo, D., Signorile, A., Papa, S., Capitanio, N., Pacelli, C. & Cocco, T. 2019. Increased Levels of cAMP by the Calcium-Dependent Activation of Soluble Adenylyl Cyclase in Parkin-Mutant Fibroblasts. Cells, 8.

63. Tavares, L. P., Negreiros-Lima, G. L., Lima, K. M., Pmr, E. S., Pinho, V., Teixeira, M. M. & Sousa, L. P. 2020. Blame the signaling: Role of cAMP for the resolution of inflammation. Pharmacol Res, 159, 105030.

64. Thakur, N., Ray, A. P., Sharp, L., Jin, B., Duong, A., Pour, N. G., Obeng, S., Wijesekara, A. V., Gao, Z. G., Mccurdy, C. R., Jacobson, K. A., Lyman, E. & Eddy, M. T. 2023. Anionic phospholipids control mechanisms of GPCR-G protein recognition. Nat Commun, 14, 794.

65. Venkatakrishnan, A. J., Deupi, X., Lebon, G., Tate, C. G., Schertler, G. F. & Babu, M. M. 2013. Molecular signatures of G-protein-coupled receptors. Nature, 494, 185–94.

66. Viedma-Poyatos, A., González-Jiménez, P., Langlois, O., Company-marín, I., Spickett, C. M. & Pérez-Sala, D. 2021. Protein Lipoxidation: Basic Concepts and Emerging Roles. Antioxidants, 10.

67. Vigor, C., Bertrand-Michel, J., Pinot, E., Oger, C., Vercauteren, J., LE Faouder, P., Galano, J. M., Lee, J. C. & Durand, T. 2014. Non-enzymatic lipid oxidation products in biological systems: assessment of the metabolites from polyunsaturated fatty acids. J Chromatogr B Analyt Technol Biomed Life Sci, 964, 65–78.

68. Wang, X., Lv, S., Sun, J., Zhang, M., Zhang, L., Sun, Y., Zhao, Z., Wang, D., Zhao, X. & Zhang, J. 2022. Caffeine reduces oxidative stress to protect against hyperoxia-induced lung injury via the adenosine A2A receptor/cAMP/PKA/Src/ERK1/2/p38MAPK pathway. Redox Rep, 27, 270–278.

69. Wang, X., Zhang, T., Song, Z., Li, L., Zhang, X., Liu, J., Liu, X., Qiu, L., Qian, Z., Zhou, S., Feng, L., Hu, G., Meng, B., Zhai, Q., Ren, X., Fu, K., Li, L., Wang, P. & Zhang, H. 2019. Tumor CD73/A2aR adenosine immunosuppressive axis and tumor-infiltrating lymphocytes in diffuse large B-cell lymphoma: correlations with clinicopathological characteristics and clinical outcome. Int J Cancer, 145, 1414–1422.

70. Weis, W. I. & Kobilka, B. K. 2018. The Molecular Basis of G Protein-Coupled Receptor Activation. Annu Rev Biochem, 87, 897–919.

71. string-name>Welihinda, A. A., Kaur, M., Greene, K., Zhai, Y. & Amento, E. P. 2016. The adenosine metabolite inosine is a functional agonist of the adenosine A2A receptor with a unique signaling bias. Cell Signal, 28, 552–60.

71. Xu, F., Wu, H., Katritch, V., Han, G. W., Jacobson, K. A., Gao, Z. G., Cherezov, V. & Stevens, R. C. 2011. Structure of an agonist-bound human A2A adenosine receptor. Science, 332, 322–7.

72. Yan, L., Burbiel, J. C., Maass, A. & Muller, C. E. 2003. Adenosine receptor agonists: from basic medicinal chemistry to clinical development. Expert Opin Emerg Drugs, 8, 537–76.

73. Ye, L., Van Eps, N., Zimmer, M., Ernst, O. P. & Prosser, R. S. 2016. Activation of the A2A adenosine G-protein-coupled receptor by conformational selection. Nature, 533, 265–8.

74. Zhang, B., Li, S. & Shui, W. 2022. Post-Translational Modifications of G Protein-Coupled Receptors Revealed by Proteomics and Structural Biology. Front Chem, 10, 843502.

