## Supplementary for "The effect of oxidative stress on the Adenosine A2a Receptor structure, activity and signalling" (from another version of this preprint: version 1, 2024-10-31)

### Supplementary Figure 1

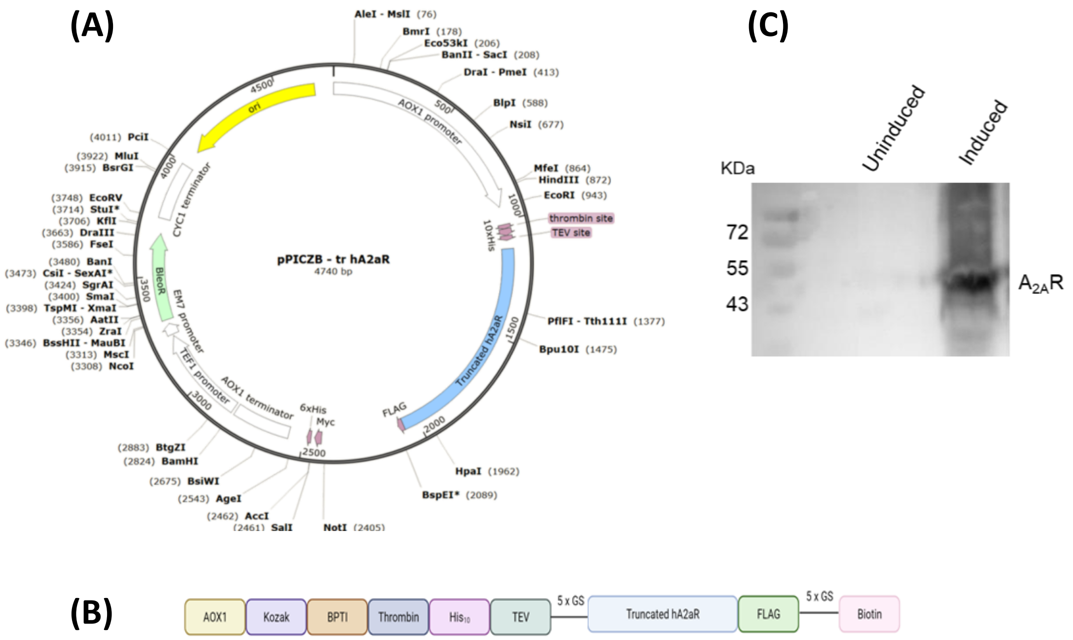


**Expression of the A2aR in *Pichia pastoris***. (A) Multi-tagged de-glycosylated C-terminal truncated A2AR (A316) cloned into a pPICZB plasmid. (B) Linear sequence showing the location of all tags. (C) The A2aR was expressed in P. pastoris SMD1163 cells stably transformed with the pPICZB-A2aR plasmid and induced by growth in the presence of methanol. The expression was assessed by SDS-PAGE and Western blotting using a specific primary antibody against the A2aR and a horseradish peroxidase (HRP)-conjugated secondary antibody.

**Supplementary Figure 2**

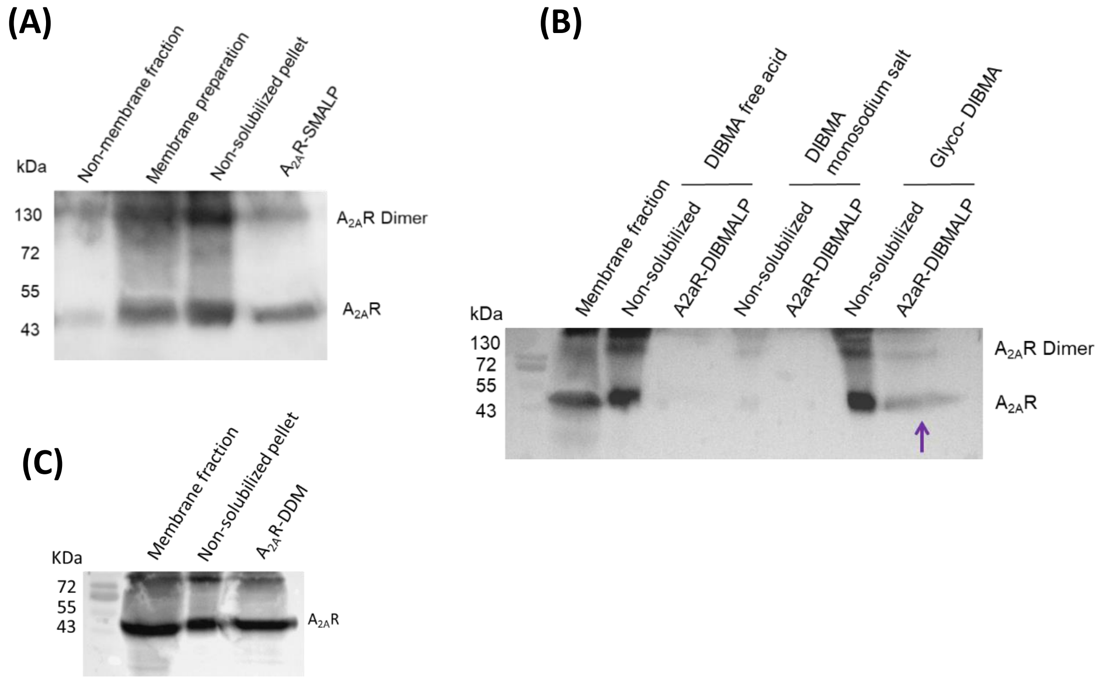


**A2aR solubilization from Pichia pastoris cell membranes**. The A2aR was solubilized from the membrane preparation using SMA2000 at 2.5% (w/v) (A); DIBMA free acid, DIBMA monosodium salt, and Glyco-DIBMA at a final concentration of 5% (B); or the detergent DDM (C). Aliquots of the non-membrane fraction, membrane preparation, non-solubilized material and solubilized protein were analysed by SDS-PAGE and Western blotting using an anti-A2aR primary antibody and HRP-conjugated secondary antibody.

**Supplementary Figure 3**

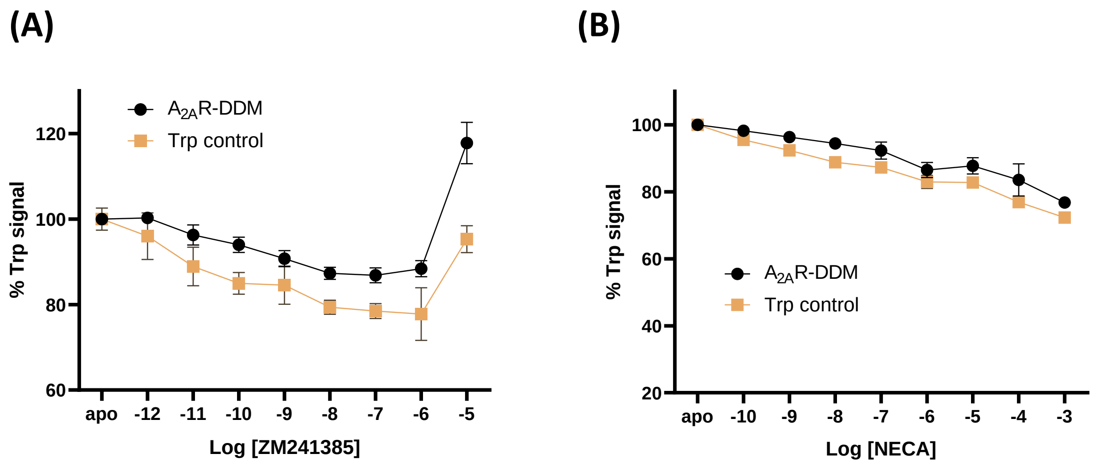


**Ligand-induced changes in A2aR-DDM fluorescence.** The intensity of fluorescence emission of A2AR-SMALP before (apo) and after the addition of increasing concentrations of (A) the antagonist ZM241385 and (B) the agonist NECA. Results are presented as a percentage of the summed intensity of fluorescence emission (290-500 nm) relative to apo A2aR-SMALP or initial Trp control as 100%. The Trp control is included to determine non-specific effects of the buffers or ligands interacting with Trp and affecting the fluorescence. Data are presented as the mean of all replicates ± SD (n=3).
It can be seen that the DDM-A2aR appeared different to the trp control, but the decrease in fluorescence observed was opposite to that observed with SMALP-A2aR.

**Supplementary Figure 4**

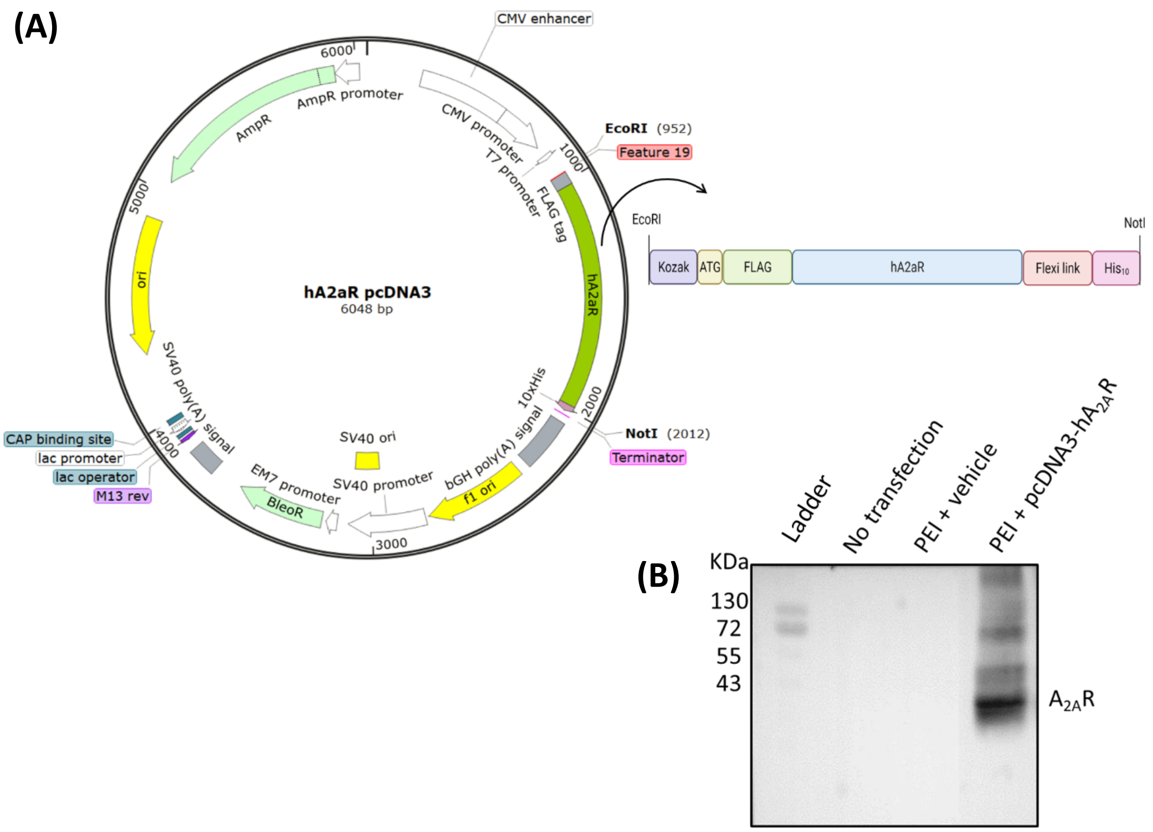


**Design of the A2AR-pcDNA3 plasmid and detection of A2aR expression**. (A) Schematic representation of the A2AR insert and final construct of the A2AR cloned into the pcDNA3 vector. In design of the plasmid, all the available sequences of the hA2AR in the membrane protein database "Membrane proteins of Known 3D structure" (mpstruc, Stephen White laboratory at UC Irvine and the Bridge Institute at USC) that had been employed to study the crystal structure of the A2AR were utilized. The FASTA sequences of all relevant structures were obtained from the Protein Data Bank (PDB, 177910 Biological Macromolecular Structures). The endogenous sequence of the human A2A Receptor published in UniprotKB - P29274 (AA2AR_HUMAN) (2002 – 2021 UniProt Consortium) was obtained as a FASTA file and included in the comparison. FASTA sequences were clustered using the multiple sequence alignment tool "Clustal Omega" from EMBL-EBI (EMBL-EBI, Genome Campus, Hinxton, Cambridgeshire, CB10 1SD, UK to obtain the amino acid sequence. The reverse translation server "Sequence Manipulation Suite: Reverse translate" (2000, 2004 Paul Stothard) was used to obtain the nucleotide sequence. FLAG tag (DYKDDDDK), deca-histidine tag, Kozak consensus sequence (AT), unstructured flexible linker (GGSGSG) and EcoRI and NotI restriction sites were added to the sequence. The designed sequence was purchased from Eurofins Scientific cloned into a pEX-K248 standard vector. (B) HEK293T cells seeded onto 6-well plates were transiently transfected with 2 µg of pcDNA3-hA2AR clone 3 and 8 µL polyethylenimine (PEI). A non-transfected and a vehicle-transfected control with 1X PBS were included in this experiment. Protein expression was demonstrated in cell lysates analysed by western blotting using a specific anti-A2AR primary antibody and HRP-conjugated secondary antibody.

**Supplementary Table 1**. Unique peptides identifying the A2aR protein by LC-MS/MS.

| Identified protein | Form | Digestion | Matches | Unique peptides | Peptide score |
|---|---|---|---|---|---|
| AA2AR_HUMAN | Monomer | Trypsin | 2 | KQMESQPLPGER | 54 |
| | | | | RYNGLVTGTR | 43 |
| AA2AR_HUMAN | Dimer | Chymotrypsin | 3 | LAIAIDRYI | 40 |
| | | | | FEDVVPIMNY | 40 |
| | | | | LSHTNSVVNPF | 40 |
| AA2AR_HUMAN | Monomer | Chymotrypsin | 7 | LNSNLQNVTNY | 61 |
| | | | | AAADIAVGVL | 58 |
| | | | | AITISTGF | 43 |
| | | | | SLLAAIDRY | 61 |
| | | | | AIGLTPMLGW | 43 |
| | | | | FEDVVPIMNY | 60 |
| | | | | SHTNSVVNPFIY | 72 |

**Supplementary Table 2**. Identification of the A2AR expressed in mammalian HEK293T cells by in-gel digestion and LC-MS/MS.

| Enzyme | Taxonomy | Protein | Description | Score | Unique peptides |
|---|---|---|---|---|---|
| **Trypsin** | Homo sapiens | A2AR_HUMAN | Adenosine receptor A2a OS=Homo sapiens OX=9606 GN=ADORA2A PE=1 SV=2 | 104 | YNGLVTGTR [a] QMESQPLPGER [b] |
| **Chymo-trypsin** | Homo sapiens | A2AR_HUMAN | Adenosine receptor A2a OS=Homo sapiens OX=9606 GN=ADORA2A PE=1 SV=2 | 49 | SHTNSVVNPF [c] |

[a] located between intracellular loop 2 (ICL2) and transmembrane helix 4 (TM4)
[b] located in intracellular loop 3 (ICL3) between transmembrane helices 5 (TM5) and 6 (TM6)
[c] located in the central region of the transmembrane helix 7 (TM7).
